# Identification of actionable targeted protein degradation effector sites through Site-specific Ligand Incorporation-induced Proximity (SLIP)

**DOI:** 10.1101/2025.02.04.636303

**Authors:** Zhangping Xiao, Efthymios Spyridon Gavriil, Fangyuan Cao, Xinyue Zhang, Stan Xiaogang Li, Sergei Kotelnikov, Patrycja Michalska, Friederike Marte, Chloe Huang, Yudi Lu, Yunxuan Zhang, Erika Bernardini, Dima Kozakov, Edward W. Tate

## Abstract

Targeted protein degradation (TPD) is a rapidly emerging and potentially transformative therapeutic modality. However, the large majority of >600 known ubiquitin ligases have yet to be exploited as TPD effectors by proteolysis-targeting chimeras (PROTACs) or molecular glue degraders (MGDs). We report here a chemical–genetic platform, Site-specific Ligand Incorporation-induced Proximity (SLIP), to identify actionable (“PROTACable”) sites on any potential effector protein in intact cells. SLIP uses genetic code expansion (GCE) to encode copper-free “click” ligation at a specific effector site in intact cells, enabling *in situ* formation of a covalent PROTAC-effector conjugate against a target protein of interest (POI). Modification at actionable effector sites drives degradation of the targeted protein, establishing the potential of these sites for TPD. Using SLIP, we systematically screened dozens of sites across E3 ligases and E2 enzymes from diverse classes, identifying multiple novel potentially PROTACable effector sites which are competent for TPD. SLIP adds a powerful approach to the proximity-induced pharmacology (PIP) toolbox, enabling future effector ligand discovery to fully enable TPD, and other emerging PIP modalities.

## INTRODUCTION

Chemically induced proximity (CIP) is a powerful concept in chemical biology and drug discovery whereby a bifunctional molecule or molecular glue brings two or more biomolecules into close spatial proximity. CIP between suitable effectors and target proteins may lead to diverse outcomes including degradation, stabilization, inhibition or activation of the targeted protein.^1,2^ In addition to providing novel tools for chemical biology, CIP also holds tremendous potential to address intractable drug targets such as transcription factors, which lack well-defined functional sites for traditional ligand binding.^3^ Targeted protein degradation (TPD) has emerged as the most successful application of CIP to date, in which a monovalent molecular glue degrader (MGD) or bivalent conjugate such as a proteolysis targeting chimera (PROTAC) mediates a ternary complex between a degradation pathway effector protein and a protein of interest (POI), directing the latter to degradation.^4^ MGDs typically facilitate a cooperative protein-protein interaction (PPI),^5^ and rational design of MGD is still challenging due to the complexity of the ternary complex interface.^6,7^ In contrast, a bifunctional PROTAC can be designed by linking discrete ligands to a POI and a component of the ubiquitin-proteasome system (UPS), typically an E3 ubiquitin ligase, thereby catalysing POI ubiquitination and degradation.^8,9^ In the past decade, more than 400 disease-related POIs have been targeted by PROTACs,^10^ and over 30 TPD drug candidates are under clinical trials against diseases from cancers to neurodegenerative conditions.^11^

Whilst the range of POIs has expanded rapidly, <2% of known E3 ligases have been exploited for TPD with just two, cereblon (CRBN) and Von Hippel–Lindau (VHL), accounting for the large majority of examples thanks to widespread expression and availability of suitable ligands.^12^ Identification of novel effectors thus offers the potential to dramatically broaden the scope of TPD,^13^ mitigating the risk of resistance to CRBN- or VHL-based PROTACs,^14^ and achieving selectivity through tissue- or disease-specific ligase expression.^12^ However, the complex ubiquitin ligase cascade is extensively regulated, and progress has been hindered by a lack of information on which specific ligandable site leads to effective POI degradation. Ubiquitination does not necessarily result in protein degradation, with K48-linked polyubiquitination most likely to lead to TPD.^15^ Furthermore, whilst POI presentation to a canonical E3 substrate recognition surface is more likely to induce effective TPD, such surfaces are poorly defined for the vast majority of ligases. Prior to investing substantial resources for rational discovery of an effective ligand for TPD, it would be ideal to pinpoint the optimal ligand binding site on a “PROTACable” E3 ligase, or other UPS component. To date, several groups have applied high throughput protein domain tagging, for example through Halo or FKBP12 fusions, resulting in the identification of many putative TPD effectors (Figure 1a).^16,17^ However, fusion of an entire domain is typically restricted to the N- or C-terminus, can alter the structure, function and localization of an effector,^18^ and provides no direct evidence for pharmacological recruitment of an effector through CIP.^16,17^ Therefore, a method to positively identify PROTACable sites on effectors in high throughput could greatly facilitate informed effector ligand discovery and subsequent TPD drug development.

**Figure 1.**
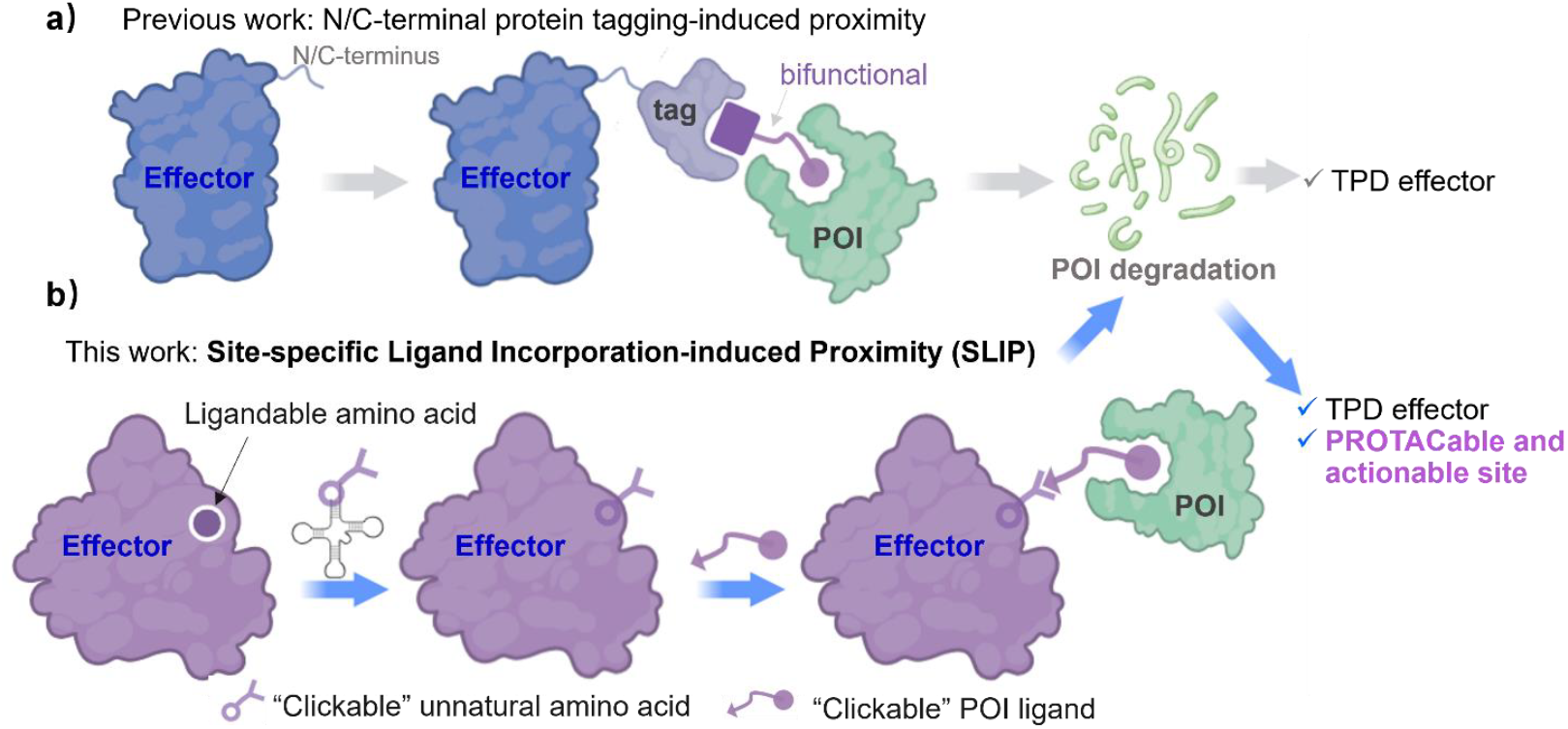
SLIP enables direct prioritization of specific effector sites for targeted protein degradation (TPD). (a) A ligand binding domain (“tag”) such as Halo or FKPB12 is fused to the N- or C-terminus of an effector to enable ligand-induced proximity to a protein of interest (POI) through a conserved motif; (b) SLIP employs genetic code expansion (GCE) to incorporate an unnatural amino acid (UAA), followed by an in-cell copper-free click reaction to precisely induce proximity at an actionable site with minimal disruption to effector structure or function. SLIP enables the identification of specific PROTACable effector sites. This figure was generated using BioRender.

Targeted covalent ligands react with nucleophilic residues such as cysteine on a target protein, and have become enormously popular in modern drug discovery due to their potential to confer higher potency and extended duration of action.^19–21^ Targeted covalency at an effector protein is also emerging as powerful approach for both MGDs and PROTACs, reducing the complexity of TPD to “pseudo-binary” kinetics while retaining a catalytic mechanism.^22^ Examples include ligands developed for a pre-selected E3 ligase,^23–26^ or through phenotypic screening and target deconvolution.^27,28^ Prioritization of ligandable nucleophilic residues presents an attractive option to simplify effector engagement and ligand discovery, but very little is known about which residues present actionable binding sites to support the rational TPD effector ligand discovery.

In the present study, we present the first systematic approach for high-throughput discovery of novel PROTACable sites to inform future TPD effector ligand discovery: Site-specific Ligand Incorporation-induced Proximity (SLIP). SLIP uses genetic code expansion (GCE) technology to incorporate an unnatural amino acid (UAA) which permits in-cell ligation to a linker and a POI ligand at any desired site on a potential effector. SLIP-mediated modification at a PROTACable site causes degradation of the targeted protein (Figure 1b), enabling direct prioritization of actionable effector sites for rational ligand development and future TPD drug discovery.

## RESULTS AND DISCUSSION

### Site-specific ligand incorporation through genetic code expansion (GCE)

To enable precise E3 ligase modification in cells with minimal perturbation of structure and function genetic code expansion was employed to site specifically incorporate an unnatural amino acid (UAA), facilitating subsequent intracellular bioconjugation through bioorthogonal “click” chemistry.^29–31^ The widely-used *Methanosarcinales mazei* pyrrolysyl-tRNA synthetase/pyrrolysyl-tRNA (Pyl-RS/tRNA) pair was selected to install a UAA bearing a constrained ring capable of catalyst free biorthogonal ligation in intact cells, via reassignment of an *amber* TAG stop codon.^31,32^ To test this system an all-in-one plasmid was designed encoding a double mutant PylRS with expanded UAA recognition, PylRS[Y306A/Y384F], alongside Pyl tRNA and E3 ligase VHL (Figure 2a). An internal ribosome entry site (IRES) motif was used to decouple VHL translation from Pyl-RS, which was in turn linked via a ribosomal skipping peptide (P2A) to enhanced green fluorescent protein (GFP), to facilitate selection of transfected cells.

**Figure 2.**
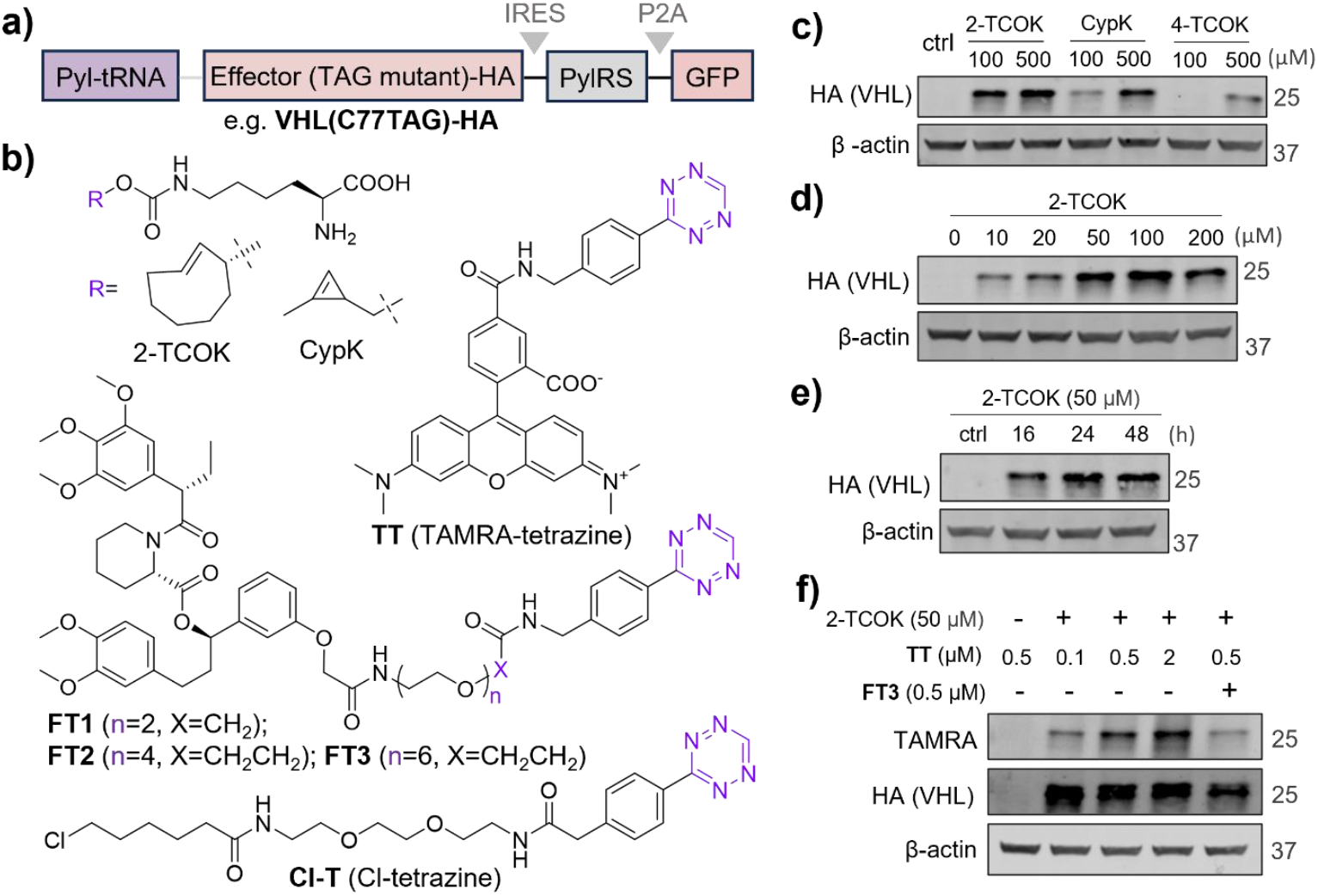
Site-specific ligand incorporation via genetic code expansion. (a) All-in-one plasmid encoding the orthogonal Pyl-RS/tRNA pair and VHL_Cys77TAG-HA for expression of UAA-modified VHL. GFP is used to monitor transfection efficiency and analysis. (b) Structures of UAAs and tetrazine derivatives used to perform Cu-free “click” ligations in cells. (c) Immunoblot analysis of protein expression efficiency with different UAA supplements; cells were transfected and treated with DMSO or different UAA for 30 hours, and expression analyzed by anti-HA Western blot, with β-actin as loading control. (d) 2-TCOK dose-dependency study; cells were transfected and incubated with different concentrations of 2-TCOK for 30 hours, and expression analyzed by anti-HA Western blot. (e) Time-dependence of VHL_Cys77TAG-HA expression with addition of 50 µM 2-TCOK, analyzed by anti-HA Western blot. (f) In-gel fluorescence analysis of 2-TCOK substituted protein labelling in live cells; cells were transfected and incubated with or without 50 µM 2-TCOK for 30 hours, then washed, pretreated with DMSO or 500 nM **FT3** for 30 min before labeling with TAMRA-tetrazine (**TT**) at the concentration indicated for 1 hour, and cell lysates analyzed by in-gel fluorescence and anti-HA Western blot.

For the first proof of concept we focused on VHL Cys77, generating Cys77 to TAG mutant VHL_C77TAG. Prior evidence suggests that this site may be both ligandable through covalency,^33^ and PROTACable based on the observation that conjugation of a bromodomain ligand via a maleimide linker at VHL Cys77 induced degradation of BRD4 following electroporation of recombinant purified conjugate into cells.^34^ To evaluate intracellular expression of VHL_C77TAG under GCE, HEK293 cells were transfected with an all-in-one plasmid encoding hemagglutinin (HA) tagged protein (VHL_C77TAG-HA), using an optimized transfection method (Figure S1). These cells were incubated with different lysine derivatives linked via a carbamate to a strained alkene, 2- or 4-substituted *trans*-cyclooctene (2-/4-TCOK) or methyl cyclopropene (CypK), capable of bioorthogonal ligation to a tetrazine derivative via inverse electron demand Diels-Alder (IEDDA) cycloaddition (Figure 2b). Anti-HA Western blot demonstrated that 100 µM 2-TCOK afforded clear VHL_C77TAG-HA UAA-dependent expression (Figure 2c) in line with the known substrate preference of PylRS-AF[Y306A/Y384F],^31^ whereas CypK and 4-TCOK showed lower efficiency. Hypothesizing that excess UAA may be difficult to wash out from cells and can potentially compete with ligand-modified VHL, we selected 2-TCOK for further studies to optimize VHL_C77TAG-HA expression dependence on UAA concentration and time. VHL_C77TAG-HA was dependent on the presence of 2-TCOK, with detectable expression at concentrations as low as 10 µM at 30 hours incubation, increasing to a maximum at 50 µM (Figure 2d). Optimization of incubation time at 50 µM 2-TCOK identified maximum expression at 24 hours onwards (Figure 2e).

TAMRA-tetrazine (**TT**, Figure 2b) was used to probe the efficiency of in-cell bioorthogonal ligation to 2-TCOK-substituted VHL_C77TAG-HA by in-gel fluorescence. As expected, fluorescent labelling of VHL_C77TAG-HA was only observed in the presence of 2-TCOK, and exhibited dose-dependency from 0.1 to 2 µM **TT**, with more unspecific labelling at 2 µM **TT** (Figure 2f). Labelling was markedly reduced by pretreating cells with 0.5 µM **FT3** (Figure 2b), a non-fluorescent probe designed to enable induction of proximity (see below), indicating cellular uptake of **FT3** and modification in competition with **TT**. These data demonstrate that tetrazine-tagged ligands can be conjugated to 2-TCOK-substituted VHL under optimized conditions in intact cells, encouraging us to move on to study targeted degradation using the SLIP platform.

### Targeted protein degradation through site-specific POI ligand incorporation

We next set out to study the effect of POI ligand incorporation using FKBP12^F36V^-mCherry protein fusion as a model system.^35^ A cell line stably expressing FKBP12^F36V^-mCherry was generated through transduction with a lentiviral construct followed by single-cell clone selection by fluorescence-activated cell sorting (FACS) (Figure S2). As expected, both dTAG^V^-1^36^ (VHL-based PROTAC) and dTAG-13^35^ (CRBN-based PROTAC) potently induced mCherry degradation in this cell line (Figure S3) as measured by FKBP12^F36V^-mCherry fluorescence intensity, establishing a quantitative readout for PROTAC-induced TPD of this construct. We hypothesized that site-specific bioorthogonal ligation of an FKBP12^F36V^ ligand to VHL would facilitate FKBP12^F36V^-mCherry degradation in a similar manner by bringing it into proximity (Figure 3a), if the ligand is conjugated at a PROTACable site.

**Figure 3.**
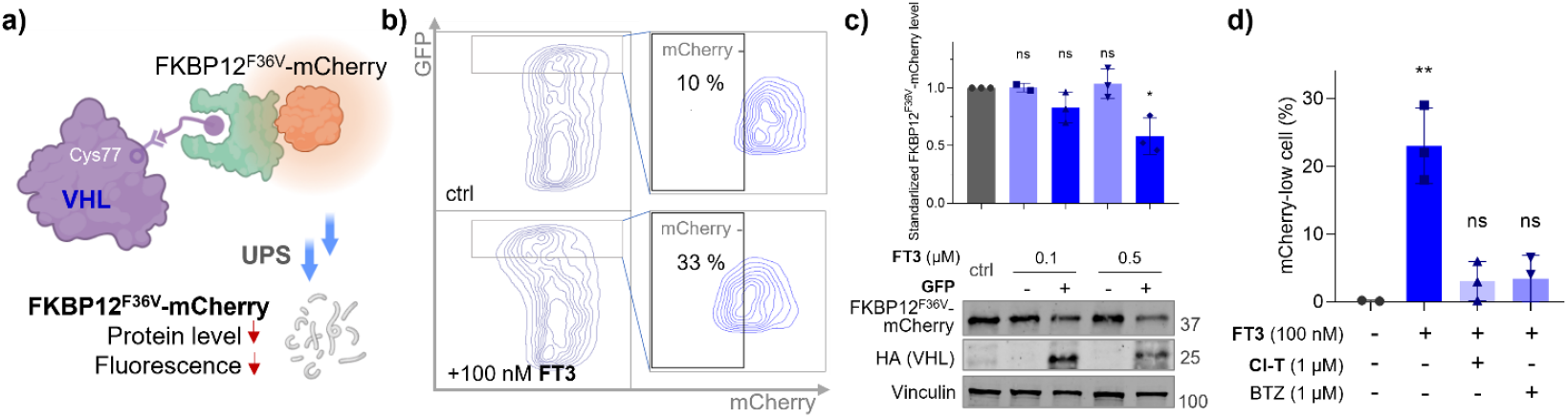
Site-specific FKBP12^F36V^ ligand incorporation induces TPD of FKBP12^F36V^-mCherry protein. (a) Schematic hypothesis for FKBP12^F36V^ ligand-bioconjugation induced proximity and subsequent protein degradation. FKBP12^F36V^-mCherry degradation can be assessed by fluorescent signal reduction. (b) HEK293 cells stably expressing FKBP12^F36V^-mCherry were transfected with plasmid containing VHL_C77TAG in the presence of 50 µM 2-TCOK for 30 hours, washed, treated with DMSO or **FT3** (100 nM) for 16 hours, and analyzed by flow cytometry. (c) HEK293 cells stably expressing FKBP12^F36V^-mCherry were transfected with VHL_C77TAG in the presence of 50 µM 2-TCOK for 30 hours, washed, treated with DMSO or **FT3** (100 or 500 nM) for 16 hours, and sorted by GFP expression using FACS between the top 10% (GFP-high) and the bottom 90% (GFP-low); FKBP12^F36V^-mCherry levels were then analyzed by immunoblot detected by anti-mCherry antibody, which was normalized to loading control Vinculin. (d) FKBP12^F36V^-mCherry expressing cells were transfected to express VHL_C77TAG-HA in the presence of 50 µM 2-TCOK for 30 hours, washed, pre-treated with Cl-T or Bortezomib (BTZ) for one hour, then with **FT3** (100 nM) for 16 hours, followed by cytometry analysis. Values are shown as means ± SD (n=3, \**p*<0.05 and \*\**p*<0.01).

To test this hypothesis we transiently expressed VHL_C77TAG in the presence of 2-TCOK and treated cells with either DMSO or **FT3** to conjugate a FKBP12^F36V^ ligand to position 77 of VHL via a hexapolyethylglycol (PEG6) linker, using copper-free IEDDA ligation. FKBP12^F36V^-mCherry degradation was monitored by fluorescence flow cytometry, revealing a population of cells shifted to a lower mCherry signal (mCherry-low) following 100 nM **FT3** treatment, compared to DMSO-treated control (Figure 3b). In GFP-high cells which most strongly express the all-in-one construct and therefore likely also 2-TCOK substituted VHL, 33% mCherry-low signal was observed following **FT3** treatment compared to a baseline of 10% in DMSO-treated cells, consistent with ligand-incorporation induced FKBP12^F36V^-mCherry degradation. Separating GFP-high (top 10%) and GFP-low (bottom 90%) cells by FACS enabled direct confirmation of **FT3**-dependent protein level decrease in FKBP^F36V^-mCherry (Figure 3c). About 40% FKBP^F36V^-mCherry protein level decrease was observed in GFP-high cells upon 500 nM **FT3** treatment. Taken together, these data suggest that site-specific FKBP12^F36V^ ligand incorporation through 2-TOCK at VHL position 77 leads to FKBP12^F36V^-mCherry protein level decrease.

Next, we tested the hypothesis that ligand-modified VHL induces FKBP12^F36V^-mCherry through a targeted protein degradation mechanism, whereby ligand conjugation leads to induced proximity through a binary complex, followed by VHL-mediated proteasome-dependent destruction of the target protein. Pretreatment with excess tetrazine analogue lacking the FKBP12^F36V^ binding moiety (**Cl-T**) to block all potential ligation sites, or with proteosome inhibitor bortezomib, efficiently rescued mCherry signal loss on subsequent treatment with **FT3**, consistent with the requirement for VHL ligand conjugation and proteasome-dependent degradation (Figure 3d). Finally, we demonstrated that swapping VHL for an unrelated protein lacking E3 ligase activity, nanoluciferase, caused no change in mCherry signal across two different 2-TCOK modified mutants (nanoluciferase N35TAG and K55TAG) under the same conditions of **FT3** treatment (Figure S5). Taken together, these data are consistent with targeted degradation of FKBP12^F36V^-mCherry driven by Site-specific Ligand Incorporation-induced Proximity (SLIP) to active VHL, followed by proteosome-mediated degradation.

### SLIP differentiates PROTACable sites on VHL

VHL is a substrate recognition subunit of the Cullin 2 RING E3 ligase complex, in which VHL forms an interface with adaptor proteins ElonginC and ElonginB (VCB-Cul2).^37^ Hypoxia-inducible factor (HIF), a principle physiological VCB-Cul2 substrate, engages the VHL binding site through a hydroxylated proline residue, a critical interaction which is mimicked by VHL-recruiting PROTACs.^38^ We hypothesized that the SLIP platform can be used to probe the potential PROTACability of varied positions on VHL to identify actionable sites for future effector ligand development. To this end, we generated VHL mutants at the canonical substrate recognition site (H110TAG) and a distal site at the ElonginC/VHL interface C162TAG, complementing C77TAG which is located between these sites when mapped onto a published X-ray structure of a PROTAC-bound BRD4-VCB complex (Figure 4a).^37^ All three HA-tagged variants were expressed in cells at a similar level in the presence of 2-TCOK, although at a level ca. >20-fold lower than wild-type VHL-HA (Figure 4b). **FT3** incorporation induced notable mCherry reduction for C77TAG and H110TAG, but C162TAG remained unchanged (Figure 4c). Furthermore, a hook effect was observed using **FT3** concentrations ranging from 20 nM to 2 µM on both C77 and H110, consistent with a PROTAC-like mechanism of action. Maximal mCherry reduction was achieved at 200 nM **FT3**, and markedly diminished at 2 µM **FT3**, indicating competition for binding POI by excess **FT3** above that required to fully engage 2-TCOK modified VHL.

**Figure 4.**
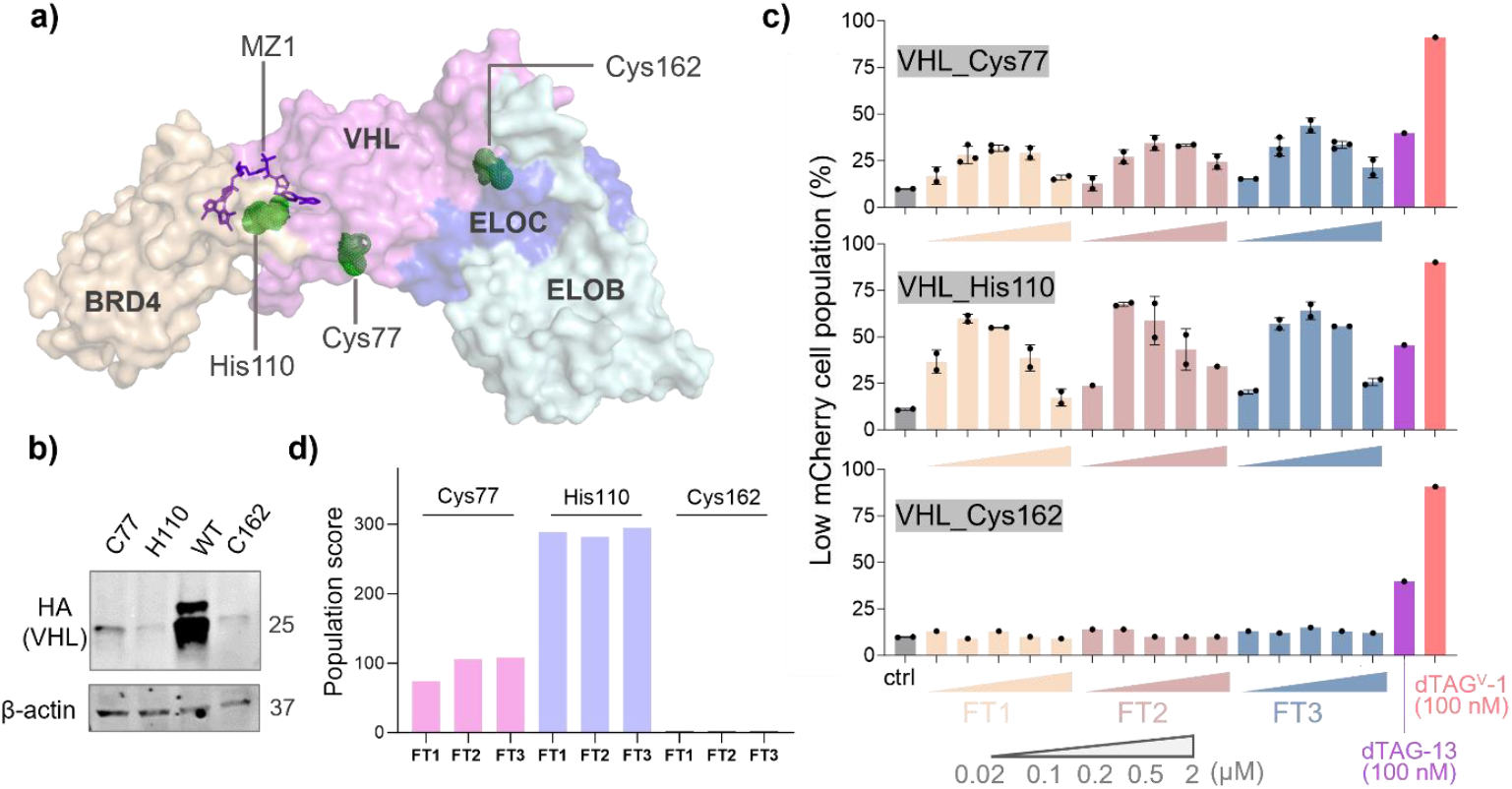
SLIP enables combinatorial and differential interrogation of PROTACable sites and linker chemistry. (a) Structure of ternary complex BRD4-MZ1-VCB (VHL and adaptor proteins Elongin C and Elongin B), mutated-sites highlighted in green (PDB: 5T35^37^). (b) HEK293 cells were transiently transfected with wild-type or different variants of VHL in the presence of 50 µM 2-TCOK for 30 hours, VHL protein levels was accessed by immunoblot using anti-HA antibody. (c) HEK293 cells stably expressing FKBP12^F36V^-mCherry were transfected with different VHL variants in the presence of 50 µM 2-TCOK for 30 hours, washed, treated with DMSO (ctrl), dTAG PROTACs, or different concentrations of tetrazine probes for 16 hours, as indicated; mCherry reduction was analyzed by flow cytometry as described above. (d) Population of generated feasible low-energy complex conformations formed between ligand-modified VCB and FKBP12^F36V^. The computational studies were performed using the fast Fourier transform (FFT)-based PROTAC complex modeling method.^42^

SLIP also enables exploration of the relationship between linker length and degradation potency. Three FKBP12^F36V^ ligand analogues containing different linkers to tetrazine (PEG2, PEG4, or PEG6, Figure 2a) were tested at five different concentrations from 20 nM to 2 µM for each of the three VHL variants, alongside wild type VHL control. Interestingly, PEG6-linked **FT3** performed optimally at C77TAG, delivering 34% mCherry signal reduction, with decreasing efficiency at decreasing chain length (PEG4 to PEG2), indicating that a longer linker is most effective at the Cys77 site. All three probes demonstrated comparable potency at H110TAG, and in all cases a distinct hook effect was observed, reaching a maximum mCherry degradation of 68%. In contrast, no degradation was induced for any probe at any tested concentration for C162TAG, or for overexpression of wild-type VHL (Figure S6). In comparison, a PROTAC engaging the canonical VHL substrate site (dTAG^V^-1) delivered FKBP12^F36V^-mCherry degradation somewhat inferior to SLIP at H110TAG, resulting in 80% signal reduction at 100 nM PROTAC, whilst a CRBN-engaging PROTAC (dTAG-13) delivered 36% reduction after the same exposure time. These findings demonstrate that SLIP can assess both the PROTACability of different sites on an effector protein and dependency on linker length, enabling systematic exploration of linker chemistry (also termed “linkerology”^39^).

To understand the structural basis for PROTACability at each site, we employed fast Fourier transform (FFT)-based PROTAC complex modeling method to find energetically favorable poses of complexes formed between ligand-modified VHL and FKBP12^F36V^.^40,41^ We found that the number of feasible low-energy complex conformations for different probes and sites aligned well with optimal POI degradation potency (Figure 4d, Figure S7), increasing from 79 conformations for **FT1** at position 77 to 101 and 102 for **FT2** and **FT3** at the same site, respectively. Approximately 300 conformations were found for **FT1, FT2**, or **FT3** at position 110, whilst no favorable conformation could be generated for position 162. These results are consistent with efficient TPD driven by propensity to form a binary complex, suggesting that SLIP selects suitable sites for future PROTAC development.

### SLIP identifies new potentially PROTACable sites across varied TPD effector proteins

We next turned to proof of concept for extension of SLIP across a library of potential PROTACable sites across diverse potential TPD effectors, focusing primarily on known covalently ligandable cysteines as a privileged class of actionable sites for covalent PROTAC or MGD discovery.^43^ We combined information from several ligandable cysteine databases^33,44,45^ and findings from previous studies on PROTACable E3 and E2 ligases to prioritize a set of 22 sites on 17 potential effectors (Figure S8).^16,17^

The E3 ligase CRBN is the most widely exploited effector for TPD, leading to marketed drugs in the immunomodulatory imide drug (IMiD) class which act as MGDs at the well-studied imide binding site,^6^ as well as the large majority of PROTACs in clinical development.^8^ Novel sites enabling recruitment of CRBN for TPD are highly sought after since they may prevent the undesired degradation activities of IMiDs, which may lead to toxicity or teratogenicity and limit the applications of IMiD-derived PROTACs or MGDs, particularly for indications beyond oncology.^14^ CRBN His352 resides at the imide binding site, and has been successfully exploited by a covalent PROTAC bearing a fluorosulfate IMiD analogue, providing a validated site to apply SLIP.^23^ We replaced His352 with 2-TCOK and confirmed that this site is also potently PROTACable by SLIP, featuring PROTAC-like dependency on both linker length and **FT** probe concentration, with a maximum potency 1.3-fold that of VHL_C77TAG (Figure 5a).^23^ Three cysteines reported to be ligandable out of 16 total across the CRBN sequence were selected for further investigation, at positions 218, 286 and 393.^16^ Cys218 was identified as a promising site for POI degradation, reaching more than half the efficiency of reference variant VHL_C77TAG. In each case, activity displayed a hook effect with respect to probe concentration. However, Cys286 or Cys393 induced no detectable degradation which may be explained by the relatively buried nature of Cys286 and the involvement of Cys393 in zinc binding (Figure 5b). These results support the generality of SLIP for identification of PROTACable sites on an effector beyond VHL, including Cys218 as a promising alternative binding site on CRBN for future covalent PROTAC discovery.

**Figure 5.**
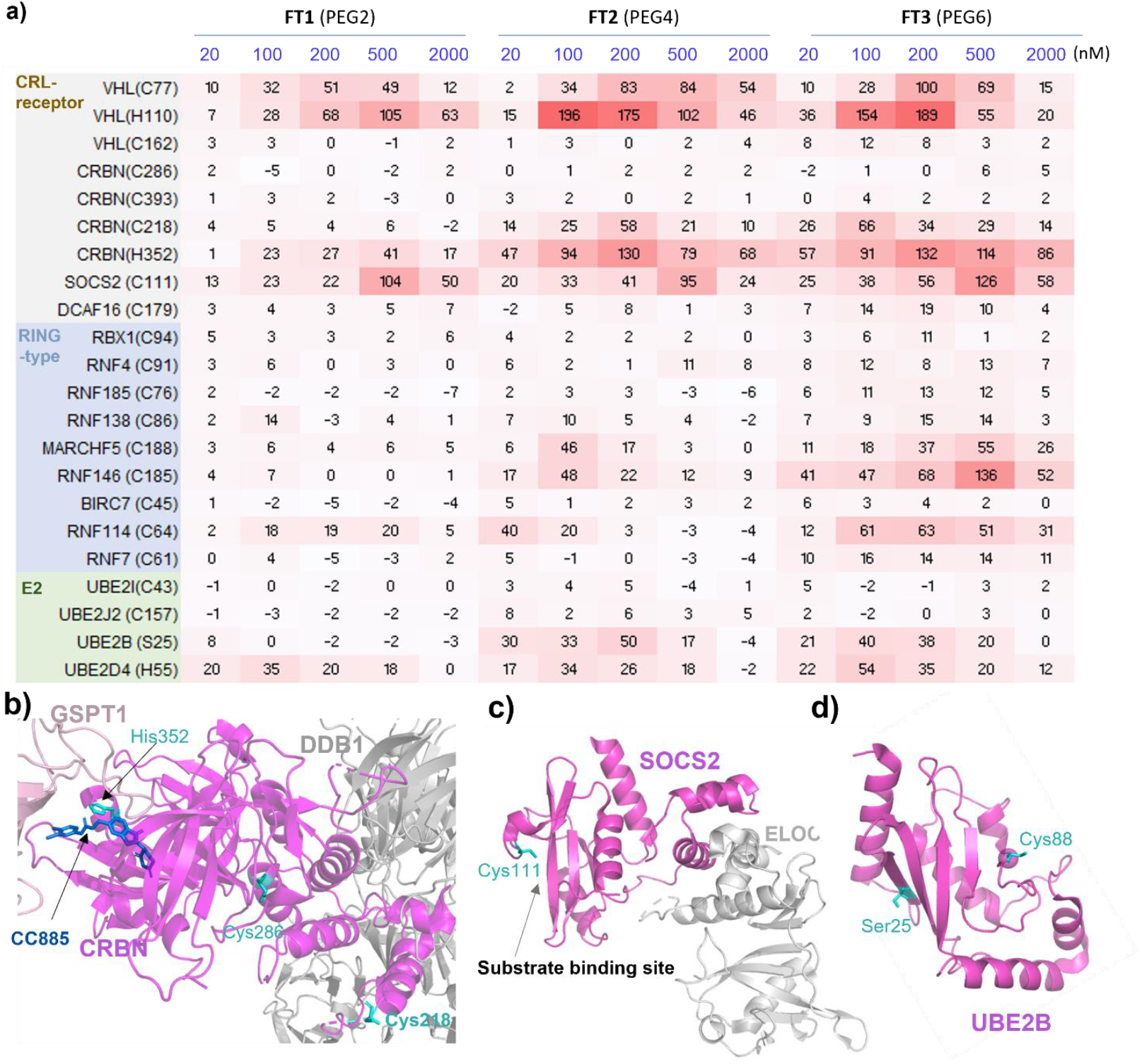
Broad identification of PROTACable sites and effectors using SLIP. (a) FKBP12^F36V^-mCherry expressing cells were transfected with different variants in the presence of 50 µM 2-TCOK for 30 hours, washed, treated with different concentrations of tetrazines for 16 hours, and mCherry level analyzed via flow cytometry normalized to the maximal effect of VHL_C77TAG. (b) Structure of GSPT1/CC-885/CRBN/DDB1 complex (PDB: 5HXB^50^). (c) Structure of SOCS2-Elongin C/B complex (PDB: 2C9W^51^). (d) Structure of UBE2B (PDB: 2YB6^52^). Figures were prepared using PyMOL. Effector proteins are shown in magenta, and key residues are highlighted in cyan.

To demonstrate the general applicability of SLIP, a set of 11 reported ligandable cysteines were selected on 16 different E3 ligases, most of which had some prior evidence using generic domain fusions supporting potential recruitment for TPD but not to date exploited by ligands.^16^ Out of these sites, SOCS2 Cys111 and RNF146 Cys185 were found to be particularly efficient, exhibiting even higher potency than the benchmark variant VHL_C77TAG. SOCS2 Cys111 is located at the E3 ligase substrate binding site, encouraging future development of PROTACs based on reported covalent ligands targeting this cysteine (Figure 5c).^46^ RNF146 is a RING finger protein with limited information regarding functional surfaces, but these results suggest that Cys185 may be located on or close to the substrate recognition site. In addition, we identified position MARCHF5 Cys188 and RNF114 Cys64 as promising PROTACable sites, with maximal degradation about half that of VHL_Cys77.

Whilst E2 conjugating enzymes typically act as the catalytic core for ubiquitination and are recruited by E3 ligases to induce substrate ubiquitination, they also have the potential to act directly as TPD effectors.^47,48^ Therefore, we extended SLIP to identify PROTACable sites on E2 enzymes, investigating four sites on four E2 ligases with previous identified as potential effectors by protein domain tagging.^16^ Since E2 enzymes must be charged with ubiquitin by E1 ligases at an active site cysteine and selectively targeting cysteines outside this site may prove challenging,^49^ we expanded our analysis beyond cysteines to potentially ligandable residues such as serine and histidine.^23,24^ Interestingly, UBE2B Ser25 and UBE2D4 His55 delivered half the activity of VHL_C77TAG, with contrasting optimal linker lengths: **FT2** (PEG4) for UBE2B Ser25 and **FT3** (PEG6) for UBE2D4 His55. Ser25 is located on the face opposite to the UBE2B catalytic residue Cys88 (Figure 5d), suggesting that engaging Ser25 may pose minimal risk of interfering with UBE2B activity. Two other sites, UBE2J2 Cys151 and UBE2I Cys43, showed no degradation activity in line with data from domain tagging which suggests that UBE2B and UBE2D4 are more potent potential degraders than UBE2J2 and UBE2I.^16^

### Endogenous protein degradation by SLIP

We next explored the versatility of SLIP to target endogenous proteins for degradation. CRBN-recruiting PROTACs have been successfully applied to TPD of the kinase Aurora A,^53,54^ and we used tetrazine-linked Aurora A probes **AT1** and **AT2** to modify CRBN via 2-TCOK substitution at His352 (Figure 6a). Following FACS separation of cells transfected with CRBN_H325TAG all-in-one plasmid into high and low GFP populations, immunoblot analysis showed that **AT1** induced Aurora A protein level decreased in CRBN_H325TAG high expressing cells at 500 nM and 1 µM **AT1**, relative to CRBN_H325TAG-low cells (Figure 6b). The PROTACable hits SOCS2_C111 and RNF146_C185 from our screening were also evaluated in this experiment. Cells expressing either SOCS2 mutant or RNF146 mutant showed decreased Aurora A protein level after 500 nM **AT1** or **AT2** treatment (Figure 6c). These results demonstrated that the SLIP system can induce endogenous protein degradation, establishing Cys111 on SOCS2 and Cys185 on RNF185 as highly promising PROTACable sites for novel PROTACs discovery.

**Figure 6.**
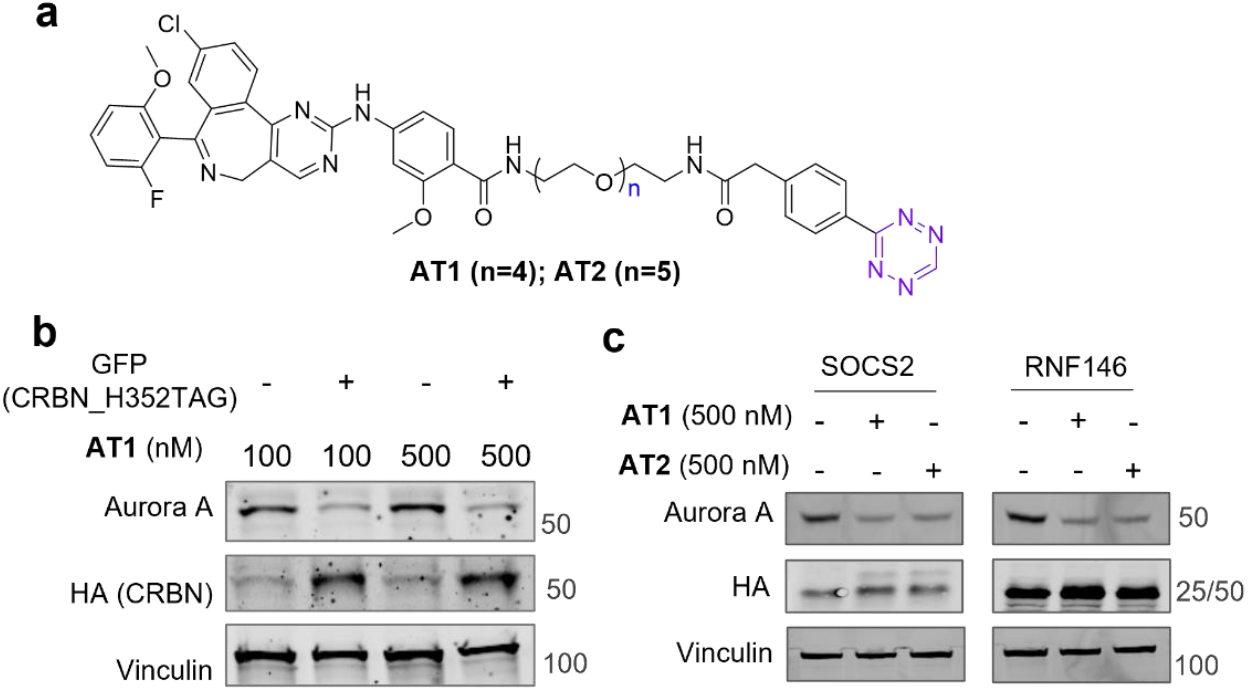
SLIP induces targeted degradation of endogenous Aurora A protein. (a) Structures of tetrazine-modified Aurora A ligand **AT1** and **AT2**. (b) Cells were transfected with CRBN_H325TAG all-in-one plasmid (see Figure 2a) for 30 hours in the presence of 50 µM 2-TCOK, washed, treated with 71 µM cycloheximide (CHX, which blocks additional protein synthesis) and **AT1** at the concentration indicated for 4 hours, separated by FACS into GFP high and low populations, and Aurora A protein levels analyzed by immunoblot following cell lysis. (c) Following the same protocol as in (b), cells were transfected with either SOCS2_C111TAG or RNF146_C185TAG, treating with **AT1** or **AT2** for 4 hours.

## CONCLUSION

Chemically induced proximity (CIP) is emerging as an effective and precise approach for the study of protein function, and to support therapeutic discovery. Current methods to systematically probe the potential of novel CIP effectors typically involve fusing the protein under study to a ligand binding domain such as FKBP or Halo and have delivered significant insights.^35,55,56^ However, these domains are >14 kDa and only straightforwardly introduced at the N- or C-terminus, with potential drawbacks including alterations to protein function or localization, steric interference with physiological or neo-substrates, and dissociation of the CIP site of action from plausible ligand binding sites, limiting the transition from potential effector to pharmacological proof of concept.^18^

Here we introduce a conceptually complimentary approach which precisely induces proximity between a potential effector and a POI in cells. SLIP enables assessment of the PROTACability of TPD effector proteins with single amino acid level precision, requiring minimal modification to the TPD effector. The additional mass introduced by GCE is ca. 100-fold smaller than that of a protein domain fusion and can be introduced at any desired position in any effector with only a single codon change, enabling systematic proteome-wide screening for actionable sites. GCE results in modified effector expression ca. 10-fold lower than wild type which may offer an advantage in gain-of-function studies, which can be particularly susceptible to false positive artefacts driven by overexpression.^57^ Furthermore, SLIP establishes a direct connection between a potential covalent or non-covalent ligand binding site and propensity to induce TPD for both model substrates and endogenous proteins for which a suitable ligand exists. It also enables combinatorial screening of effector sites against diverse POI ligand and linker chemistry, with interesting trends emerging even from the small set of chain lengths explored here. In combination with efficient and high accuracy computational modeling of induced proximity such as FFT based PROTAC modeling approach,^42^ SLIP may provide valuable information to assist PROTAC linker design.

SLIP recapitulates potent TPD at known PROTACable and substrate binding sites for VHL and CRBN, and we anticipate that it will facilitate rapid prioritization of ligand discovery at high-value actionable sites and effectors for development of PROTACs and MGDs. It may also offer a useful tool for discovery of physiological functional interfaces of effectors, given the correlation between substrate binding sites and PROTACability. VHL Cys77 and CRBN Cys218 were identified as promising distal PROTACable sites, offering alternative approaches to engage these potent TPD effectors, overcome drug resistance to current PROTACs or MGDs, or mitigate undesired neosubstrate degradation by IMiDs.^14^ SLIP also identified a set of potential PROTACable sites on effectors which have not yet been applied in TPD. SOCS2 Cys111 is a promising site for PROTAC ligand development based on a recently reported covalent ligand for this cysteine.^46^ Other novel sites include RNF146 Cys185, MARCHF5 Cys188, RNF114 Cys64, UBE2B Ser25 and UBE2D4 His55, offering unprecedented opportunities for *de novo* TPD effector ligand discovery and degrader development and the potential for targeted pharmacology based on tissue-specific effector expression; for example, RNF114 is known to have high and specific expression in human testis.^58^ SLIP could also be applied to screen actionable sites on a POI, although additional care may need to be taken to ensure that GCE does not influence the inherent degradability of the POI. Indeed, during preparation of the present manuscript, a similar approach targeting a POI was reported very recently,^59^ further supporting the broad applicability of this concept.

Finally, thanks to the versatility of GCE, we envisage that SLIP will be applicable in principle to any effector class with a measurable phenotype arising from proximity-induced pharmacology. Future applications for this platform beyond TPD would include protein stabilization through deubiquitinase (DUB) recruitment, protein (de)phosphorylation via kinases or phosphatases, and many others.^2,60^ SLIP may prove particularly valuable for proof of concept for new CIP concepts where a ligand is unavailable, or where current approaches are ineffective. The effectors involved in modalities beyond TPD typically exhibit more stringent specificity than ubiquitin ligases^2,61,62^ and may require more proximal and precise orientation of substrate and effector than can be achieved by a protein fusion.

## ASSOCIATED COTENT

The supporting information is available free of charge. Supporting information includes detailed chemical synthesis and characterization, details and characterization of molecular cloning and cell line generation, experimental details and additional results on cellular experiments, and computational modeling details (PDF).

## Author Contributions

E.W.T. carried out conceptualization. E.W.T., D.K. and Z.X. were responsible for funding acquisition. Z.X., E.S.G., F.C., X.Z., S.X.L., S.K., P.M., F.M., C.H., Y.L., Y.Z. and E.B. carried out the investigation. Z.X., E.S.G. and P.M. developed the methodology. S.X.L. and S.K. performed the computational work. E.W.T. and Z.X. managed project administration. Z.X. and E.W.T. performed data visualization.

E.W.T. and D.K. provided supervision. Z.X. and E.W.T. were responsible for writing the original draft. All authors have given approval to the final version of the manuscript.

## Note

The authors declare the following competing financial interest(s): EWT is or has been employed as a consultant or scientific advisory board member for Myricx Pharma, Samsara Therapeutics, Roche, Novartis and Fastbase; research in his group has been funded by Pfizer Ltd, Kura Oncology, Daiichi Sankyo, Oxstem, Exscientia, Myricx Pharma, AstraZeneca, Vertex Pharmaceuticals, GSK and ADC Technologies. EWT holds equity in Myricx Pharma, Exactmer and Samsara Therapeutics, and is a named inventor on patents filed by Myricx Pharma, Exactmer, Imperial College London and the Francis Crick Institute.

## ACKNOWLEDGMENT

We wish to acknowledge the support of the LMS/NIHR Flow Facility at Imperial for the experiments involving cell sorting (FACS). We thank Dr. Anne Schuhmacher, Dr. Jack Houghton, Dr. Mahesh Mohan and Matthew White for useful discussions. We thank Daria Berezina and Dr. Edward Bartlett for their discussion and support on cell line generation and molecular cloning.

## Table of Contents

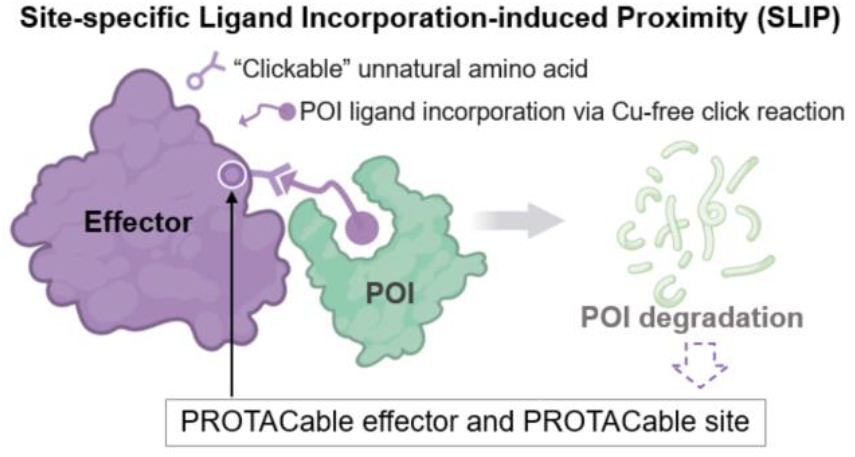

## REFERENCES

(1) Stanton, B. Z.; Chory, E. J.; Gerald R. Crabtree. Chemically Induced Proximity in Biology and Medicine. Science (80-.). 2018, 359 (6380), eaao5902.

(2) Liu, X.; Ciulli, A. Proximity-Based Modalities for Biology and Medicine. ACS Cent. Sci. 2023, 9 (7), 1269–1284.

(3) Popow, J.; Farnaby, W.; Gollner, A.; Kofink, C.; Fischer, G.; Wurm, M.; Zollman, D.; Wijaya, A.; Mischerikow, N.; Hasenoehrl, C.; et al. Targeting Cancer with Small-Molecule Pan-KRAS Degraders. Science 2024, 385 (6715), 1338–1347.

(4) Tsai, J. M.; Nowak, R. P.; Ebert, B. L.; Fischer, E. S. Targeted Protein Degradation: From Mechanisms to Clinic. Nat. Rev. Mol. Cell Biol. 2024, 25 (9), 740–757.

(5) Schreiber, S. L. Molecular Glues and Bifunctional Compounds: Therapeutic Modalities Based on Induced Proximity. Cell Chem. Biol. 2024, 31 (6), 1050–1063.

(6) Fischer, E. S.; Böhm, K.; Lydeard, J. R.; Yang, H.; Stadler, M. B.; Cavadini, S.; Nagel, J.; Serluca, F.; Acker, V.; Lingaraju, G. M.; et al. Structure of the DDB1-CRBN E3 Ubiquitin Ligase in Complex with Thalidomide. Nature 2014, 512 (1), 49–53.

(7) Schreiber, S. L. The Rise of Molecular Glues. Cell 2021, 184 (1), 3–9.

(8) Békés, M.; Langley, D. R.; Crews, C. M. PROTAC Targeted Protein Degraders: The Past Is Prologue. Nat. Rev. Drug Discov. 2022, 21 (3), 181–200.

(9) Cao, C.; He, M.; Wang, L.; He, Y.; Rao, Y. Chemistries of Bifunctional PROTAC Degraders. Chem. Soc. Rev. 2022, 51 (16), 7066–7114.

(10) Ge, J.; Li, S.; Weng, G.; Wang, H.; Fang, M.; Sun, H.; Deng, Y.; Hsieh, C.-Y.; Li, D.; Hou, T. PROTAC-DB 3.0: An Updated Database of PROTACs with Extended Pharmacokinetic Parameters. Nucleic Acids Res. 2024, 2127, 1–6.

(11) Zhong, G.; Chang, X.; Xie, W.; Zhou, X. Targeted Protein Degradation: Advances in Drug Discovery and Clinical Practice. Signal Transduct. Target. Ther. 2024, 9 (1), 308.

(12) Liu, Y.; Yang, J.; Wang, T.; Luo, M.; Chen, Y.; Chen, C.; Ronai, Z.; Zhou, Y.; Ruppin, E.; Han, L. Expanding PROTACtable Genome Universe of E3 Ligases. Nat. Commun. 2023, 14 (1).

(13) Schneider, M.; Radoux, C. J.; Hercules, A.; Ochoa, D.; Dunham, I.; Zalmas, L. P.; Hessler, G.; Ruf, S.; Shanmugasundaram, V.; Hann, M. M.; et al. The PROTACtable Genome. Nat. Rev. Drug Discov. 2021, 20 (10), 789–797.

(14) Hanzl, A.; Casement, R.; Imrichova, H.; Hughes, S. J.; Barone, E.; Testa, A.; Bauer, S.; Wright, J.; Brand, M.; Ciulli, A.; et al. Functional E3 Ligase Hotspots and Resistance Mechanisms to Small-Molecule Degraders. Nat. Chem. Biol. 2023, 19 (3), 323–333.

(15) Swatek, K. N.; Komander, D. Ubiquitin Modifications. Cell Res. 2016, 26 (4), 399–422.

(16) Poirson, J.; Cho, H.; Dhillon, A.; Haider, S.; Imrit, A. Z.; Lam, M. H. Y.; Alerasool, N.; Lacoste, J.; Mizan, L.; Wong, C.; et al. Proteome-Scale Discovery of Protein Degradation and Stabilization Effectors. Nature 2024, 628 (8009), 878–886.

(17) Serebrenik, Y. V.; Mani, D.; Maujean, T.; Burslem, G. M.; Shalem, O. Pooled Endogenous Protein Tagging and Recruitment for Systematic Profiling of Protein Function. Cell Genomics 2024, 100651.

(18) Vandemoortele, G.; Eyckerman, S.; Gevaert, K. Pick a Tag and Explore the Functions of Your Pet Protein. Trends Biotechnol. 2019, 37 (10), 1078–1090.

(19) Boike, L.; Henning, N. J.; Nomura, D. K. Advances in Covalent Drug Discovery. Nat. Rev. Drug Discov. 2022, 21 (12), 881–898.

(20) Honigberg, L. A.; Smith, A. M.; Sirisawad, M.; Verner, E.; Loury, D.; Chang, B.; Li, S.; Pan, Z.; Thamm, D. H.; Miller, R. A.; et al. The Bruton Tyrosine Kinase Inhibitor PCI-32765 Blocks B-Cell Activation and Is Efficacious in Models of Autoimmune Disease and B-Cell Malignancy. Proc. Natl. Acad. Sci. U. S. A. 2010, 107 (29), 13075–13080.

(21) Lanman, B. A.; Allen, J. R.; Allen, J. G.; Amegadzie, A. K.; Ashton, K. S.; Booker, S. K.; Chen, J. J.; Chen, N.; Frohn, M. J.; Goodman, G.; et al. Discovery of a Covalent Inhibitor of KRASG12C (AMG 510) for the Treatment of Solid Tumors. J. Med. Chem. 2020, 63 (1), 52–65.

(22) Lu, D.; Yu, X.; Lin, H.; Cheng, R.; Monroy, E. Y.; Qi, X.; Wang, M. C.; Wang, J. Applications of Covalent Chemistry in Targeted Protein Degradation. Chem. Soc. Rev. 2022, 10 (9), 9243–9261.

(23) Nowak, R. P.; Ragosta, L.; Huerta, F.; Liu, H.; Ficarro, S. B.; Cruite, J. T.; Metivier, R. J.; Donovan, K. A.; Marto, J. A.; Fischer, E. S.; et al. Development of a Covalent Cereblon-Based PROTAC Employing a Fluorosulfate Warhead. RSC Chem. Biol. 2023, 4 (11), 906–912.

(24) Shah, R. R.; De Vita, E.; Sathyamurthi, P. S.; Conole, D.; Zhang, X.; Fellows, E.; Dickinson, E. R.; Fleites, C. M.; Queisser, M. A.; Harling, J. D.; et al. Structure-Guided Design and Optimization of Covalent VHL-Targeted Sulfonyl Fluoride PROTACs. J. Med. Chem. 2024, 67 (6), 4641–4654.

(25) Ward, C. C.; Kleinman, J. I.; Brittain, S. M.; Lee, P. S.; Chung, C. Y. S.; Kim, K.; Petri, Y.; Thomas, J. R.; Tallarico, J. A.; McKenna, J. M.; et al. Covalent Ligand Screening Uncovers a RNF4 E3 Ligase Recruiter for Targeted Protein Degradation Applications. ACS Chem. Biol. 2019, 14 (11), 2430–2440.

(26) Meyers, M.; Cismoski, S.; Panidapu, A.; Chie-Leon, B.; Nomura, D. K. Targeted Protein Degradation through Recruitment of the CUL4 Complex Adaptor Protein DDB1. ACS Chem. Biol. 2024, 19 (1), 58–68.

(27) Zhang, X.; Crowley, V. M.; Wucherpfennig, T. G.; Dix, M. M.; Cravatt, B. F. Electrophilic PROTACs That Degrade Nuclear Proteins by Engaging DCAF16. Nat. Chem. Biol. 2019, 15 (7), 737–746.

(28) Zhang, X.; Luukkonen, L. M.; Eissler, C. L.; Crowley, V. M.; Yamashita, Y.; Schafroth, M. A.; Kikuchi, S.; Weinstein, D. S.; Symons, K. T.; Nordin, B. E.; et al. DCAF11 Supports Targeted Protein Degradation by Electrophilic Proteolysis-Targeting Chimeras. J. Am. Chem. Soc. 2021, 143 (13), 5141–5149.

(29) Lang, K.; Davis, L.; Wallace, S.; Mahesh, M.; Cox, D. J.; Blackman, M. L.; Fox, J. M.; Chin, J. W. Genetic Encoding of Bicyclononynes and Trans-Cyclooctenes for Site-Specific Protein Labeling in Vitro and in Live Mammalian Cells via Rapid Fluorogenic Diels-Alder Reactions. J. Am. Chem. Soc. 2012, 134 (25), 10317–10320.

(30) Nikic, I.; Kang, J. H.; Girona, G. E.; Aramburu, I. V.; Lemke, E. A. Labeling Proteins on Live Mammalian Cells Using Click Chemistry. Nat. Protoc. 2015, 10 (5), 780–791.

(31) Peng, T.; Hang, H. C. Site-Specific Bioorthogonal Labeling for Fluorescence Imaging of Intracellular Proteins in Living Cells. J. Am. Chem. Soc. 2016, 138 (43), 14423–14433.

(32) Nödling, A. R.; Spear, L. A.; Williams, T. L.; Luk, L. Y. P.; Tsai, Y. H. Using Genetically Incorporated Unnatural Amino Acids to Control Protein Functions in Mammalian Cells. Essays Biochem. 2019, 63 (2), 237–266.

(33) Takahashi, M.; Chong, H. B.; Zhang, S.; Yang, T. Y.; Lazarov, M. J.; Harry, S.; Maynard, M.; Hilbert, B.; White, R. D.; Murrey, H. E.; et al. DrugMap: A Quantitative Pan-Cancer Analysis of Cysteine Ligandability. Cell 2024, 187 (10), 2536–2556.e30.

(34) Pinch, B. J.; Buckley, D. L.; Gleim, S.; Brittain, S. M.; Tandeske, L.; D’Alessandro, P. L.; Hauseman, Z. J.; Lipps, J.; Xu, L.; Harvey, E. P.; et al. A Strategy to Assess the Cellular Activity of E3 Ligase Components against Neo-Substrates Using Electrophilic Probes. Cell Chem. Biol. 2022, 29 (1), 57–66.e6.

(35) Nabet, B.; Roberts, J. M.; Buckley, D. L.; Paulk, J.; Dastjerdi, S.; Yang, A.; Leggett, A. L.; Erb, M. A.; Lawlor, M. A.; Souza, A.; et al. The DTAG System for Immediate and Target-Specific Protein Degradation. Nat. Chem. Biol. 2018, 14 (5), 431–441.

(36) Nabet, B.; Ferguson, F. M.; Seong, B. K. A.; Kuljanin, M.; Leggett, A. L.; Mohardt, M. L.; Robichaud, A.; Conway, A. S.; Buckley, D. L.; Mancias, J. D.; et al. Rapid and Direct Control of Target Protein Levels with VHL-Recruiting DTAG Molecules. Nat. Commun. 2020, 11 (1).

(37) Gadd, M. S.; Testa, A.; Lucas, X.; Chan, K. H.; Chen, W.; Lamont, D. J.; Zengerle, M.; Ciulli, A. Structural Basis of PROTAC Cooperative Recognition for Selective Protein Degradation. Nat. Chem. Biol. 2017, 13 (5), 514–521.

(38) Min, J. H.; Yang, H.; Ivan, M.; Gertler, F.; Kaelin, W. G.; Pavietich, N. P. Structure of an HIF-1α-PVHL Complex: Hydroxyproline Recognition in Signaling. Science (80-.). 2002, 296 (5574), 1886–1889.

(39) Bemis, T. A.; Clair, J. J. La;, Burkart, M. D. Unraveling the Role of Linker Design in Proteolysis Targeting Chimeras. J. Med. Chem. 2021, 64 (12), 8042–8052.

(40) Zhang, Q.; Kounde, C. S.; Mondal, M.; Greenfield, J. L.; Baker, J. R.; Kotelnikov, S.; Ignatov, M.; Tinworth, C. P.; Zhang, L.; Conole, D.; et al. Light-Mediated Multi-Target Protein Degradation Using Arylazopyrazole Photoswitchable PROTACs (AP-PROTACs). Chem. Commun. 2022, 58 (78), 10933–10936.

(41) Ignatov, M.; Jindal, A.; Kotelnikov, S.; Beglov, D.; Posternak, G.; Tang, X.; Maisonneuve, P.; Poda, G.; Batey, R. A.; Sicheri, F.; et al. High Accuracy Prediction of PROTAC Complex Structures. J. Am. Chem. Soc. 2023, 145 (13), 7123–7135.

(42) Ignatov, M.; Jindal, A.; Kotelnikov, S.; Beglov, D.; Posternak, G.; Tang, X.; Maisonneuve, P.; Poda, G.; Batey, R. A.; Sicheri, F.; et al. High Accuracy Prediction of PROTAC Complex Structures. J. Am. Chem. Soc. 2023, 145, 7123–7135.

(43) White, M. E. H.; Gil, J.; Tate, E. W. Proteome-Wide Structural Analysis Identifies Warhead- and Coverage-Specific Biases in Cysteine-Focused Chemoproteomics. Cell Chem. Biol. 2023, 30 (7), 828–838.e4.

(44) Kuljanin, M.; Mitchell, D. C.; Schweppe, D. K.; Gikandi, A. S.; Nusinow, D. P.; Bulloch, N. J.; Vinogradova, E. V.; Wilson, D. L.; Kool, E. T.; Mancias, J. D.; et al. Reimagining High-Throughput Profiling of Reactive Cysteines for Cell-Based Screening of Large Electrophile Libraries. Nat. Biotechnol. 2021, 39 (5), 630–641.

(45) Boatner, L. M.; Palafox, M. F.; Schweppe, D. K.; Backus, K. M. CysDB: A Human Cysteine Database Based on Experimental Quantitative Chemoproteomics. Cell Chem. Biol. 2023, 30 (6), 683–698.e3.

(46) Ramachandran, S.; Makukhin, N.; Haubrich, K.; Nagala, M.; Forrester, B.; Lynch, D. M.; Casement, R.; Testa, A.; Bruno, E.; Gitto, R.; et al. Structure-Based Design of a Phosphotyrosine-Masked Covalent Ligand Targeting the E3 Ligase SOCS2. Nat. Commun. 2023, 14 (1), 1–17.

(47) Taylor, J. D.; Barrett, N.; Martinez Cuesta, S.; Cassidy, K.; Pachl, F.; Dodgson, J.; Patel, R.; Eriksson, T. M.; Riley, A.; Burrell, M.; et al. Targeted Protein Degradation Using Chimeric Human E2 Ubiquitin-Conjugating Enzymes. Commun. Biol. 2024, 7 (1).

(48) Forte, N.; Dovala, D.; Hesse, M. J.; McKenna, J. M.; Tallarico, J. A.; Schirle, M.; Nomura, D. K. Targeted Protein Degradation through E2 Recruitment. ACS Chem. Biol. 2023, 18 (4), 897–904.

(49) Stewart, M. D.; Ritterhoff, T.; Klevit, R. E.; Brzovic, P. S. E2 Enzymes: More than Just Middle Men. Cell Res. 2016, 26 (4), 423–440.

(50) Matyskiela, M. E.; Lu, G.; Ito, T.; Pagarigan, B.; Lu, C. C.; Miller, K.; Fang, W.; Wang, N. Y.; Nguyen, D.; Houston, J.; et al. A Novel Cereblon Modulator Recruits GSPT1 to the CRL4 CRBN Ubiquitin Ligase. Nature 2016, 535 (7611), 252–257.

(51) Bullock, A. N.; Debreczeni, J. É.; Edwards, A. M.; Sundström, M.; Knapp, S. Crystal Structure of the SOCS2-Elongin C-Elongin B Complex Defines a Prototypical SOCS Box Ubiquitin Ligase. Proc. Natl. Acad. Sci. U. S. A. 2006, 103 (20), 7637–7642.

(52) Hibbert, R. G.; Huang, A.; Boelens, R.; Sixma, T. K. E3 Ligase Rad18 Promotes Monoubiquitination Rather than Ubiquitin Chain Formation by E2 Enzyme Rad6. Proc. Natl. Acad. Sci. U. S. A. 2011, 108 (14), 5590–5595.

(53) Adhikari, B.; Bozilovic, J.; Diebold, M.; Schwarz, J. D.; Hofstetter, J.; Schröder, M.; Wanior, M.; Narain, A.; Vogt, M.; Dudvarski Stankovic, N.; et al. PROTAC-Mediated Degradation Reveals a Non-Catalytic Function of AURORA-A Kinase. Nat. Chem. Biol. 2020, 16 (11), 1179–1188.

(54) Sells, T. B.; Chau, R.; Ecsedy, J. A.; Gershman, R. E.; Hoar, K.; Huck, J.; Janowick, D. A.; Kadambi, V. J.; Leroy, P. J.; Stirling, M.; et al. MLN8054 and Alisertib (MLN8237): Discovery of Selective Oral Aurora A Inhibitors. ACS Med. Chem. Lett. 2015, 6 (6), 630–634.

(55) Jan, M.; Scarfò, I.; Larson, R. C.; Walker, A.; Schmidts, A.; Guirguis, A. A.; Gasser, J. A.; Słabicki, M.; Bouffard, A. A.; Castano, A. P.; et al. Reversible ON- And OFF-Switch Chimeric Antigen Receptors Controlled by Lenalidomide. Sci. Transl. Med. 2021, 13 (575), 1–13.

(56) Mercer, J. A. M.; DeCarlo, S. J.; Roy Burman, S. S.; Sreekanth, V.; Nelson, A. T.; Hunkeler, M.; Chen, P. J.; Donovan, K. A.; Kokkonda, P.; Tiwari, P. K.; et al. Continuous Evolution of Compact Protein Degradation Tags Regulated by Selective Molecular Glues. Science (80-.). 2024, 383 (6688).

(57) Vetma, V.; Perez, L. C.; Eliaš, J.; Stingu, A.; Kombara, A.; Gmaschitz, T.; Braun, N.; Ciftci, T.; Dahmann, G.; Diers, E.; et al. Confounding Factors in Targeted Degradation of Short-Lived Proteins. ACS Chem. Biol. 2024, 19 (7), 1484–1494.

(58) Capon, F.; Bijlmakers, M. J.; Wolf, N.; Quaranta, M.; Huffmeier, U.; Allen, M.; Timms, K.; Abkevich, V.; Gutin, A.; Smith, R.; et al. Identification of ZNF313/RNF114 as a Novel Psoriasis Susceptibility Gene. Hum. Mol. Genet. 2008, 17 (13), 1938–1945.

(59) Shade, O.; Ryan, A.; Belsito, G.; Deiters, A. Investigating Protein Degradability through Site-Specific Ubiquitin Ligase Recruitment. RSC Chem. Biol. 2024.

(60) Peng, Y.; Liu, J.; Inuzuka, H.; Wei, W. Targeted Protein Posttranslational Modi Fi Cations by Chemically Induced Proximity for Cancer Therapy. J. Biol. Chem. 2023, 299 (4), 104572.

(61) Siriwardena, S. U.; Godage, D. N. P. M.; Shoba, V. M.; Lai, S.; Shi, M.; Wu, P.; Chaudhary, S. K.; Schreiber, S. L.; Choudhary, A. Phosphorylation-Inducing Chimeric Small Molecules. J. Am. Chem. Soc. 2020, 142, 14052–14057.

(62) Yamazoe, S.; Fu, Y.; Wu, W.; Zeng, L.; Sun, C.; Liu, Q.; Lin, J.; Lin, K.; Fairbrother, W. J.; Staben, S. T. Heterobifunctional Molecules Induce Dephosphorylation of Kinases - A Proof of Concept Study. J. Med. Chem. 2020, 63, 2807–2013.

