## Supplementary Information for "Identification of actionable targeted protein degradation effector sites through Site-specific Ligand Incorporation-induced Proximity (SLIP)"

#### Supplemental Results

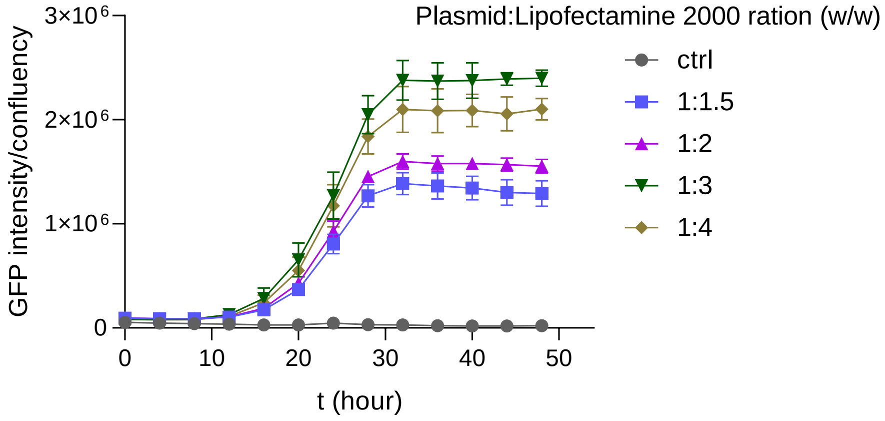

**Figure S1**. **Transient transfection condition optimization.** HEK293 cells in a 12-well plate (800 k/well) were transfected with the all-in-one plasmid with different ratio of Lipofectamine 2000. The plasmid (1.0 µg in 50 µL Opti-EM) was mixed with different amount of Lipofectamine 2000 (1.5, 2.0, 3.4, or 4.0 µg in 50 µL Opti-EM). After incubation at RT for 15 min, the mixture was added into one well of cells, followed by addition of 2-TCOK (final concentration of 50 µM) and tetracycline (final concentration of 3.3 µg/µL). The GFP expression was monitored using Incucyte S3 using green channel (Ex 441-481/ Em 503-544nm).

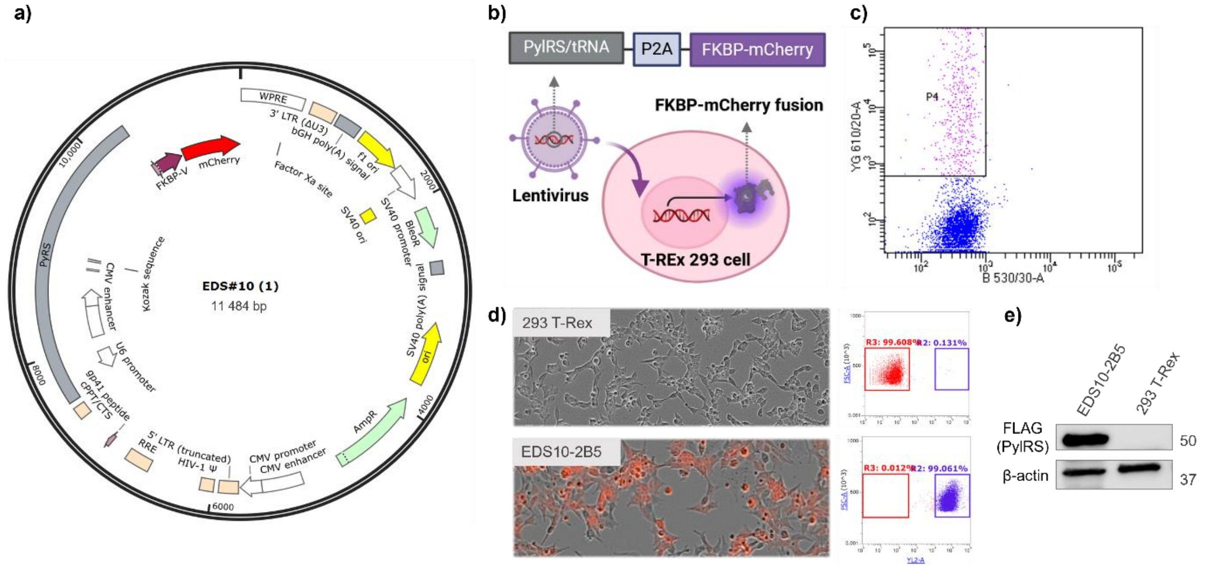

**Figure S2**. **Generation and Characterization of reporting cell line stably expressing FKBP$^{F36V}$-mCherry.** (a) The map of transfer plasmid used for lentivirus generation. This plasmid was prepared using primers listed in Table S1. (b) Schematic illustration of the mechanism of FKBP$^{F36V}$-mCherry expressing cell line generation. (c) Fluorescence-activated cell sorting (FACS) based on mCherry signaling (610/20-A channel) was used to sort mCherry-positive cells into a 96 well plate as single cells. (d) After selection under Zeocin and expansion, cells derived from the same single cell were analyzed using Incucyte S3 using red (Ex 567-607/ Em 622-704nm) fluorescent channel and flow cytometry (Attune NXT, YL2 channel, 620/15). In comparison to parental cells, EDS10-2B5 cells are homogenously mCherry positive. (e) Parental cells HEK293 T-Rex or EDS10-2B5 cells were lysed and PylRS expression was assessed using its FLAG tag. B-actin was used as a loading control.

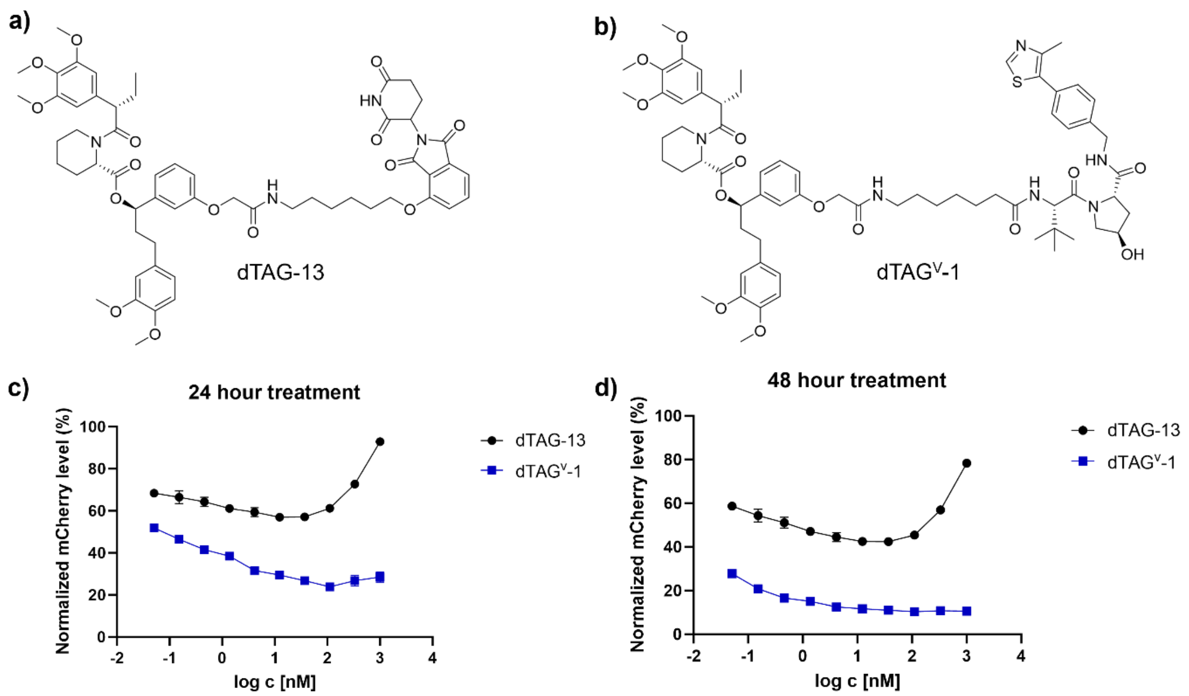

**Figure S3**. **Degradability of FKBP$^{F36V}$-mCherry.** (a) and (b) Structures of dTAG$^{V}$-1[1] (VHL-based PROTAC) and dTAG-13[2] (CRBN-based PROTAC). (c) and (d) Different concentrations of dTAG$^{V}$-1 or dTAG-13 were added to EDS10-2B5 cells in a 96-well plate (20 k/well). mCherry level was evaluated on Incucyte S3 using red (Ex 567-607/ Em 622-704nm) fluorescent channel at 24 hours and 48 hours of treatment. Red fluorescent intensity was normalised to DMSO-treated control and standardized by cell confluency (n=3).

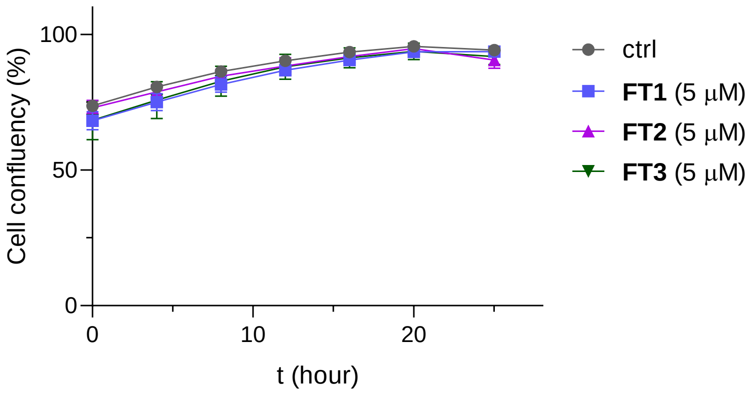

**Figure S4**. **Cytotoxicity of FKBP$^{F36V}$ ligand-tetrazines.** EDS10-2B5 cells were seeded in a 96-well plate at a density of 40 k/well. After overnight incubation, cells were treated with 5 µM FKBP$^{F36V}$ ligand-tetrazines (**FT1**, **FT2**, or **FT3**) or DMSO for 24 hours. Cell confluency was monitored on Incucyte S3 using phase object (n=3).

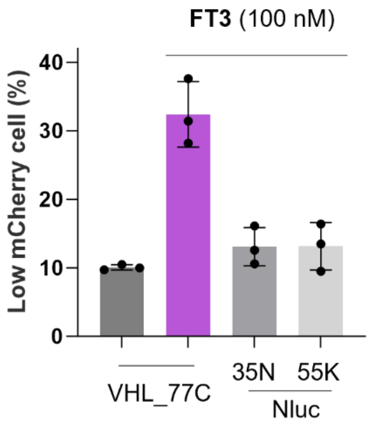

**Figure S5**. **VHL_C77TAG, but not Nluc_N35/K55TAG, causes FKBP12$^{F36V}$-mCherry protein reduction**. FKBP12$^{F36V}$-mCherry expressing cells were transfected to express VHL_C77TAG-HA or nanoluciferase (Nluc)_N35/K55TAG in the presence of 50 µM 2-TCOK for 30 hours, washed, then treated with **FT3** (100 nM) for 16 hours, followed by cytometry analysis.

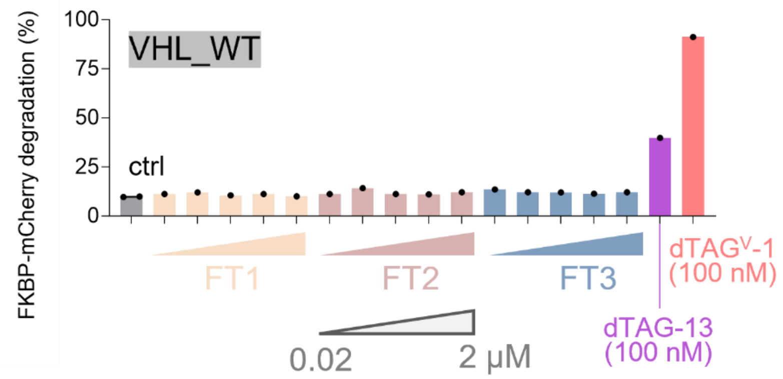

**Figure S6. VHL-HA overexpression doesn't cause FKBP12$^{F36V}$-mCherry protein degradation.** HEK293 cells stably expressing FKBP12$^{F36V}$-mCherry were transfected with different VHL variants in the presence of 50 μM 2-TCOK for 30 hours, washed, treated with DMSO (ctrl), dTAG PROTACs, or different concentrations of tetrazine probes for 16 hours, as indicated; mCherry reduction was analyzed by flow cytometry.

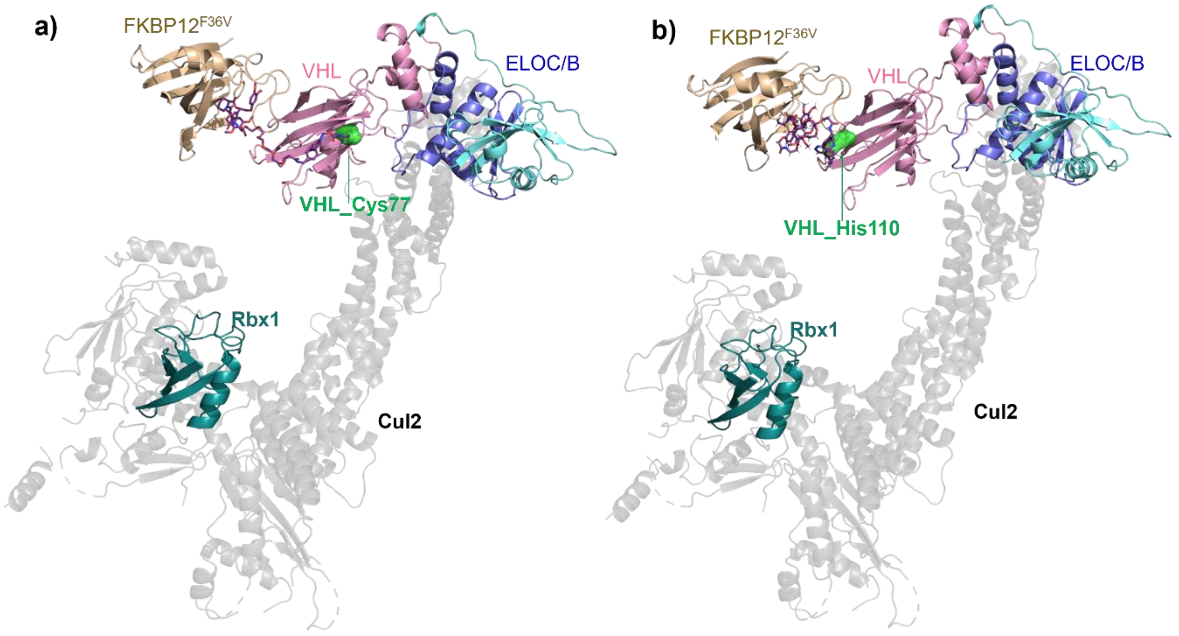

**Figure S7. Energetically favorable poses induced by ligand modifications on VHL_C77/H110.** Representative low-energy pose formed between FKBP$^{F36V}$ ligand modified VHL_C77 (a) or VHL_H110 (b) complex with FKBP$^{F36V}$. The computational studies were performed using the fast Fourier transform (FFT)-based PROTAC complex modeling method described above. The figures were prepared using PyMOL. FKBP$^{F36V}$ is shown in gold, VHL is shown in pink.

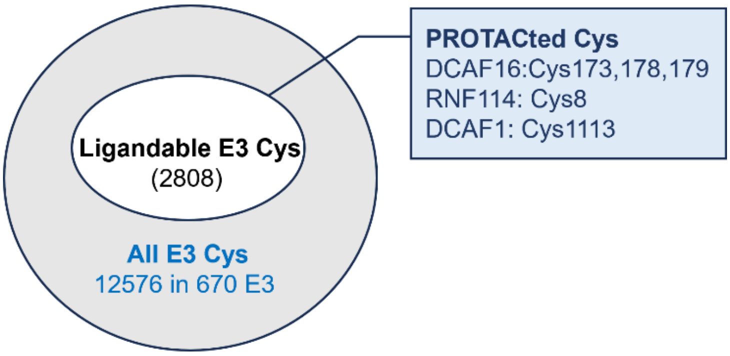

**Figure S8. Number of ligandable cysteines on E3 ligases.** The list of E3 was derived from this database (https://hanlaboratory.com/E3Atlas/)[3] with a standard of score >2 being applied. The ligandability data were mainly collected from three databases: CysDB (labelled yes)[4], SLC APBB (with CR>4)[5], Drugmap (>60% engagement)[6]. Several ligandable cysteines such as Cys8 on RNF114 have been successfully engaged in PROTAC discovery.

#### Experimental Procedures

##### Plasmids and subcloning

A complete list of vectors and primers used in this study is available in Table 1. DNA sequence amplification was conducted using Q5® Hot Start High-Fidelity DNA Polymerase kit (NEB, M0515). DNA fragments assembly was performed using the NEBuilder HiFi Assembly Kit (NEB, E5520S). Product guides were followed. NEB® 5-alpha Competent *E. coli* (NEB, C2987H) was used for transformation and single clones were selected for mini culture. Plasmids were purified using QIAprep Spin Miniprep Kit (Qiagen, 27104). Purified plasmids were sent for key segment sequencing via Sanger sequencing (AZENTA) or whole plasmid sequencing (FullCirclesLab) to confirm subcloning results. Primers used for subcloning were ordered from Thermo Fisher.

##### Cloning for all-in-one plasmid

A plasmid containing human C-terminally HA-tagged VHL and an enhanced green fluorescent protein (GFP), which are decoupled using an internal ribosome entry site (IRES) motif was ordered from GenScript. Three copies of Pyl-tRNA under a U6 promotor were inserted into the plasmid, followed by adding PylRS_FLAG to the position between IRES and GFP, linked to GFP via a self-cleaving peptide P2A. Pyl-tRNA and PylRS_FLAG were amplified from pAS_4xMma PylT_FLAG-Mma PylRS AF (Addgene, 140023).

##### Cloning for Lenti Virus plasmid

The 3rd generation Lenti virus system was used to generate stable cell line in this work. The transfer plasmid was generated using FKBP12$^{F36V}$-mCherry gene (Addgene, 72906) and PylRS_FLAG (Addgene, 140023), which were linked via a self-cleaving peptide T2A. NEB® Stable Competent *E. coli* (NEB, C3040H) was used for transformation and plasmid application.

##### Molecular cloning

##### Generation of FKBP12$^{F36V}$-mCherry expressing cell line

Lentiviral plasmid PylRS-T2A-FKBP12$^{F36V}$-mCherry, packaging and viral envelope plasmids pCMV-Delta-8.2 (Addgene, 12263) and pCMV-VSV-G (Addgene, 8454) were used to prepare lentiviral particles in HEK293-FT cells according to the instructions in the Lenti-X Lentiviral Expression System Manual (Clontech). Stable PylRS-T2A-FKBP12$^{F36V}$-mCherry expressing Flp-In™ 293 T-Rex was generated by transduction of $1 \times 10^6$ low-passage cells with lentivirus in a 6-well plate, followed by Zeocin selection at 10 μg $ml^{-1}$. After 1 week, cells were applied to prepare single cell originated monoclonal stable cell lines using fluorescence-activated cell sorting (FACS) on channel 610/20 (mCherry) into 96-well plates. Single cells were expanded and evaluated using mCherry-based flow cytometry and anti-FLAG western blot (PylRS is FLAG-tagged).

##### Mutagenesis

All mutagenesis reactions were carried out using the Site Directed Mutagenesis kit (NEB, E0554) according to the manufacturer's instructions. The list of mutations and mutagenic primer sets are shown in Table 1. All purified plasmids were sent for Sanger sequencing (AZENTA) to confirm correct mutation. Mutagenic primers were purchased from ThermoFisher.

#### Cell culture and compound preparation

HEK293T, Flp-In™ 293 T-Rex cell lines were cultured in Dulbecco's modified Eagle medium (DMEM) supplemented with GlutaMAX (Thermo Fisher, #10566016), 10% *v*/*v* FBS in a 37 °C, 5% $CO_2$ incubator. Cells were selected with 0.5 mg/mL Zeocin (Thermo Fisher, #R25001) and 10 µg/mL blasticidin S hydrochloride (Thermo Fisher, #A1113903). Tetrazine-5-TAMRA (Stratech, #CLK-017-05-JEN), Bortezomib (Themo Fisher, #J60378.MA), dTAG-13 (Merck, #SML2601), dTAG$^{V}$-1 (MedChemExpress, #HY-145514) and synthesized tetrazine analogues were prepared as 10 mM stocks in DMSO stored in -20 °C, and diluted to corresponding final concentration before use. Unnatural amino acids (UAAs) are purchased from SiChem GmbH (2-TCOK, #SC-8008; 4-TCOK, #SC-8060; CypK, #SC-8017.) and diluted in pH 8-9 PBS as 10 mM stocks and stored in -20 °C.

#### Western blot analysis

Cell lysate was prepared in RIPA buffer containing protease inhibitor cocktail (1x, Roche, C762Q77) and Benzonase (1x). Lysate (20 µg/well) was denatured with 4x Laemmli sample uuffer before being loaded onto and separated by a precast 12% SDS-PAGE gel (BioRad, #4561043). The proteins in the gel were then transferred to a nitrocellulose membrane on Trans-Blot Turbo Transfer System. After blocking with 5% (w/v) of skimmed milk for one hour at room temperature and incubation with the appropriate primary antibody diluted in TBST buffere containing 2% BSA (HA, Cell Signaling, #3956, 1:1000 dilution; β-actin, ThermoFisher, #MA1-140, 1:5000; vinculin, Abcam, #ab91459, 1:2000; Aurora A, Cell Signaling, #14475, 1:1000; mCherry, Cell Signaling, #43590, 1:1000.) overnight at 4 °C. Then the protein bands were visualized fluorescent secondary antibodies (IRDye 800CW, LICORbio, # 926-32211, 1:5000; IRDye 680RD, LICORbio, #926-68070, 1:5000.) on LICOR Odyssey and quantified with ImageJ software based on grayscale.

#### 2-TCOK modified E3 or E2 overexpression and FKBP12$^{F36V}$-mCherry degradation

A 96-well plate is coated by poly-L-lysine solution (30 µL/well), rock gently to ensure even coating of the culture surface. After 5 minutes, remove solution by aspiration and thoroughly rinse surface with sterile PBS buffer. Seed EDS10-2B5 cells in a precoated 96-well plate (40 k/well) to get 70% confluency the next day. Cells were transfected with corresponding plasmid (150 ng/well) and Lipofectamine 2000 mixture (450 ng/well) in total volume of 10 µL/well. Next, 2-TCOK (50 µM) and Tetracycline (3.3 µg/ mL) were added for a 30-hour incubation. Then the cells were washed four times with DMEM medium containing 1% FBS in two hours. Afterwards, tetrazines (**FT1**, **FT2**, and **FT3**) were diluted in washing buffer to appropriate concentrations before being applied to cells. After overnight incubation (16 hours), culture medium was removed, and cells were trypsinised for cytometry analysis. GFP and mCherry signal were recorded by BL1 channel (530/30 nm) and YL2 channel (620/15 nm) on cytometer (Attune NXT). Data were analyzed using FlowJo.

#### Structural modeling

To gain structural insight into experimentally observed degradation patterns, we performed computational modeling of PROTAC complexes using recently developed method.[7] Briefly, the method splits PROTAC in the middle of the linker, docks both PROTAC's ligands to corresponding proteins[8–10] if experimental complex structures are not available, and samples each half-PROTAC in the presence of the corresponding protein. The resulting ensembles of half-PROTAC end atom positions are projected onto the spatial docking grids. The method then performs FFT-based protein docking[11] with an additional "silent" convolution to find energetically favorable protein-protein complex poses. This "silent" term convolves the above-mentioned grids to ensure reconnection of the linker and account for PROTAC presence.[8,91211,13]The resulting structures are energetically relaxed by AMBER and clustered to produce PROTAC-mediated complex models.[14] Finally, these are filtered for ubiquitin accessibility to the target protein ($FKBP12^{F36V}$), and the population score is calculated based on the remaining models.[7]

##### UAA modified protein expression in cells

HEK293 (EDS10-2B5) cells were seeded in a 12-well plate at a density of 800k per well. After an overnight incubation, the cells were at a confluency of 70-90% and ready for transfection with corresponding plasmids containing TAG in frame genes. For transfection of 2 wells, 2 µg of plasmid was diluted into 100 µL Opti-EM, which was added to 100 µL Opti-MEM containing 6 µg Lipofectamine 2000. The mixture was incubated at RT for 15 min, 100 µL of the mixture was added into one well. Next, 2-TCOK (final concentration 50 µM) and tetracycline (final concentration 3.3 µg/ mL) were added. The treated cells were incubated for 30 hours, GFP expression was monitored using Incucyte. UAA modified protein expression was assessed via western blot using antibody against HA tag.

**Table 1. Plasmid vectors and primers used in this work.**

| Item | Description |
|---|---|
| P2A insertion_F | AggtctatataagcagagctGGATCCGGCGCAACAAAC |
| P2A insertion_R | GTCTCCACCTGCACTCCCATCGGTCCAGGATTCTCTTCG |
| Pyl-tRNA insertion_1_F | gcgatgtacgTAGGCGTTTTGCGCTGCTTC |
| Pyl-tRNA insertion_1_R | cgccatttttACCCTAAGCAGATTCTTCATGC |
| Pyl-tRNA insertion_2_F | tgcttagggtAAAAATGGCGGAAACCCC |
| Pyl-tRNA insertion_2_R | aaaacgcctaCGTACATCGCGAAGGTCG |
| PylRS insertion_1_F | tcctggaccgATGGTGTCTAAGGGCGAAG |
| PylRS insertion_1_R | ccatggtggcTTATCATCGTGTTCTTCAAAGG |
| PylRS insertion_2_F | acgatgataaGCCACCATGGACTACAAG |
| PylRS insertion_2_R | tagacaccatCGGTCCAGGATTCTCTTC |
| VHL_H110TAG_F | CAGAAGGATCtagAGCTATAGAGGACACCTG |
| VHL_H110TAG_R | CCTGTGCCAGGTGGC |
| VHL_C162TAG_F | AAGGAACGGTagCTGCAGGTGG |
| VHL_C162TAG_R | CAGGGTGTAGACTGGCAG |

#### Chemistry

##### General

All commercially available reagents and solvents were used without additional purification unless stated otherwise. Air and moisture sensitive reactions were carried out in dried suba-sealed flasks purged with nitrogen. Thin layer chromatography (TLC) was performed on aluminium backed plates coated with Merck DC Kieselgel 60 F254 and visualised under UV light at 254 nm. For liquid chromatography combined with mass spectrometry (LCMS) the Agilent Infinity II 1260 LCMS system was used. LC traces were recorded on a 1260 Infinity II Diode Array Detector HS scanning at 220 and 260 nm, and MS spectra were recorded on a LC single-quad InfinityLab LC/MSD MS either in positive or negative mode. LCMS Method: Mobile phase A - 0.1% formic acid in water; Mobile phase B - acetonitrile. Gradient 95% A – 5% B to 5% A – 95% B over 7 min. Flash column chromatography was carried out using glass chromatography columns packed with 40-63 μm silica gel or with the Biotage® Selekt high performance automated flash purification system. NMR spectra were recorded at room temperature (20 ± 1˚C) on Bruker 400 or 500 MHz. $^{1}$H NMR data is presented with the following: Chemical shifts (δ) are reported in parts per million (ppm) relative to residual solvent peaks as internal standards, multiplicity (s = singlet, d = doublet, t = triplet, q = quartet, m = multiplet, br = broad) coupling constants (*J*) in Hz, and integration. High resolution mass spectrometry (HRMS) was performed using electrospray ionization (ES) and time-of-flight (TOF) mass analysis. The purity of all final compounds was > 95%, as defined by LCMS and NMR.

For the synthesis of FKBP ligand-tetrazine derivatives, commercially available **1** was first deprotected, then underwent amide coupling with *N*-Boc-protected PEG-2, PEG-4 and PEG-6 based carboxylic acids to introduce suitable linker groups. After Boc removal, the final amide compounds were formed using the FKBP ligand AP1867 **(Scheme 1)**.

**Scheme 1.** Synthesis of FKBP ligand-tetrazine derivatives[a]

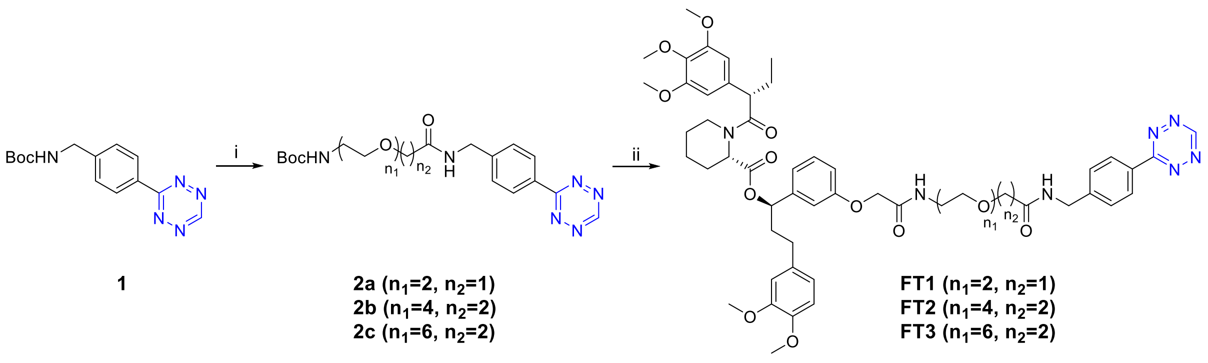

[a]Reagents and conditions. i) a. TFA, DCM, reflux,1 h; b. 2,2-dimethyl-4-oxo-3,8,11-trioxa-5-azatridecan-13-oic acid or 2,2-dimethyl-4-oxo-3,8,11,14,17-pentaoxa-5-azaicosan-20-oic acid or 2,2-dimethyl-4-oxo-3,8,11,14,17,20,23-heptaoxa-5-azahexacosan-26-oic acid, EDCI, HOBt, DCM, rt, 18 h, yield 74%-quant. over two steps. ii) a. TFA, DCM, reflux, 1 h; b. FKBP ligand AP1867, EDCI, HOBt, DCM, rt, yield 38%-quant. over two steps.

**_tert_-butyl (2-(2-(2-((4-(1,2,4,5-tetrazin-3-yl)benzyl)amino)-2-oxoethoxy)ethoxy)ethyl)carbamate (2a)**

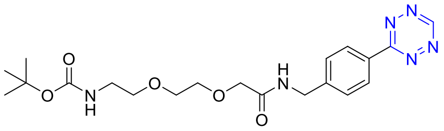

tert-butyl (4-(1,2,4,5-tetrazin-3-yl)benzyl)carbamate **1** (20 mg, 0.070 mmol) was added to a mixture of trifluoroacetic acid (2 mL) and dichloromethane (2 mL) and the reaction was refluxed for 1 h. The mixture was concentrated under reduced pressure and the crude was used in the next step without further purification. The deprotected tetrazine derivative was added to a mixture in dichloromethane (5 mL) containing 2,2-dimethyl-4-oxo-3,8,11-trioxa-5-azatridecan-13-oic acid (19 mg, 0.072 mmol), which was pre-activated by EDCI (20 mg, 0.13 mmol), HOBt (11 mg, 0.081 mmol) and triethylamine (20 uL, 0.14 mmol). The resulting mixture was stirred at room temperature for 18 h, then concentrated under reduced pressure. Purification by flash column chromatography (0-5% methanol in dichloromethane) afforded the title compound (20 mg, yield 74%) as a pink oil. $R_f$ 0.4 (5% methanol in dichloromethane). $^1$H NMR (400 MHz, $CDCl_3$) δ 10.22 (s, 1H), 8.60 (d, *J* = 8.3 Hz, 2H), 7.55 (d, *J* = 8.2 Hz, 2H), 7.42 (s, 1H), 4.73 (s, 1H), 4.63 (d, *J* = 6.2 Hz, 2H), 4.11 (s, 2H), 3.75 – 3.68 (m, 2H), 3.66 – 3.59 (m, 2H), 3.49 (t, *J* = 5.3 Hz, 2H), 3.22 (q, *J* = 5.4 Hz, 2H), 1.41 (s, 9H).

**_tert_-butyl (1-(4-(1,2,4,5-tetrazin-3-yl)phenyl)-3-oxo-6,9,12,15-tetraoxa-2-azaheptadecan-17-yl)carbamate (2b)**

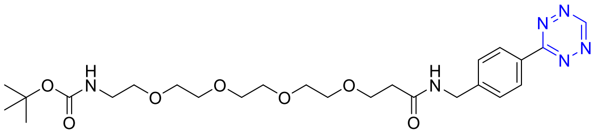

Compound **2b** was synthesized according to the procedure reported for compound **2a** using 2,2-dimethyl-4-oxo-3,8,11,14,17-pentaoxa-5-azaicosan-20-oic acid. **2b** was obtained as pink oil, yield 79%. $R_f$ 0.4 (5% methanol in dichloromethane). $^1$H NMR (400 MHz, $CDCl_3$) δ 10.20 (s, 1H), 8.57 (d, *J* = 8.2 Hz, 2H), 7.53 (d, *J* = 8.2 Hz, 2H), 7.21 (s, 1H), 5.02 (s, 1H), 4.57 (d, *J* = 6.1 Hz, 2H), 3.79 (t, *J* = 5.6 Hz, 2H), 3.67 – 3.61 (m, 4H), 3.58 – 3.53 (m, 8H), 3.48 (t, *J* = 5.1 Hz, 2H), 3.26 (q, *J* = 5.4 Hz, 2H), 2.59 (t, *J* = 5.6 Hz, 2H), 1.42 (s, 9H).

**_tert_-butyl (1-(4-(1,2,4,5-tetrazin-3-yl)phenyl)-3-oxo-6,9,12,15,18,21-hexaoxa-2-azatricosan-23-yl)carbamate (2c)**

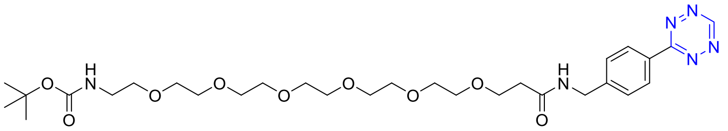

Compound **2c** was synthesized according to the procedure reported for compound **2a** using 2,2-dimethyl-4-oxo-3,8,11,14,17,20,23-heptaoxa-5-azahexacosan-26-oic acid. **2c** was obtained as pink oil in quantitative yield. $R_f$ 0.4 (5% methanol in dichloromethane). $^1$H NMR (400 MHz, $CDCl_3$) δ 10.21 (s, 1H), 8.57 (d, *J* = 8.3 Hz, 2H), 7.54 (d, *J* = 8.0 Hz, 2H), 7.26 (s, 1H), 5.09 (s, 1H), 4.58 (d, *J* = 6.1 Hz, 2H), 3.80 (t, *J* = 5.6 Hz, 2H), 3.67 – 3.55 (m, 20H), 3.52 (t, *J* = 5.1 Hz, 2H), 3.30 (q, *J* = 5.5 Hz, 2H), 2.59 (t, *J* = 5.6 Hz, 2H), 1.43 (s, 9H).

**(*R*)-1-(3-((1-(4-(1,2,4,5-tetrazin-3-yl)phenyl)-3,12-dioxo-5,8-dioxa-2,11-diazatridecan-13-yl)oxy)phenyl)-3-(3,4-dimethoxyphenyl)propyl (*S*)-1-((S)-2-(3,4,5-trimethoxyphenyl) butanoyl)piperidine-2-carboxylate (FT1)**

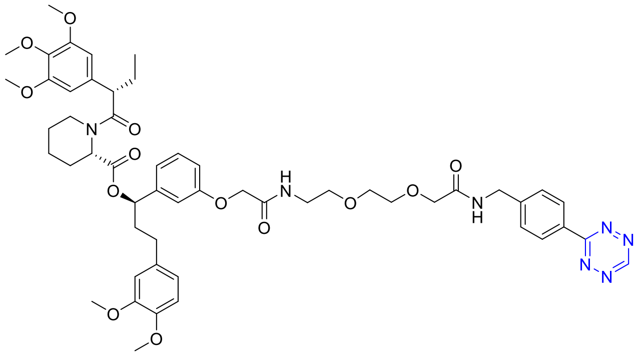

Compound **2a** (5.0 mg, 0.013 mmol) was added to a mixture of trifluoroacetic acid (2 mL) and dichloromethane (2 mL) and the reaction was refluxed for 1 h. The mixture was concentrated under reduced pressure and the crude was used in the next step without further purification. The deprotected product was added to a mixture in dichloromethane (5 mL) containing FKBP12$^{F36V}$- ligand AP1867 (5.0 mg, 0.0072 mmol), which was pre-activated by HATU (5.0 mg, 0.013 mmol) and triethylamine (50 uL, 0.35 mmol). The resulting mixture was stirred at room temperature for 18 h, then concentrated under reduced pressure. Purification by flash column chromatography (0-5% methanol in dichloromethane) afforded the title compound (7.0 mg, yield quant.) as a pink solid. $R_f$ 0.7 (10% methanol in dichloromethane). $^1$H NMR (400 MHz, $CDCl_3$) δ 10.19 (s, 1H), 8.57 (d, *J* = 10.5 Hz, 2H), 7.52 (d, *J* = 8.3 Hz, 2H), 7.44 (s, 1H), 7.16 (t, *J* = 7.9 Hz, 1H), 6.93 (t, *J* = 5.9 Hz, 1H), 6.75 (t, *J* = 9.6 Hz, 3H), 6.68 – 6.61 (m, 3H), 6.41 – 6.36 (m, 2H), 5.64 – 5.57 (m, 1H), 5.44 (d, *J* = 3.8 Hz, 1H), 4.60 (d, *J* = 6.2 Hz, 2H), 4.41 (d, *J* = 4.9 Hz, 2H), 4.09 (d, *J* = 5.3 Hz, 2H), 3.86 – 3.81 (m, 10H), 3.77 (d, *J* = 2.4 Hz, 3H), 3.69 – 3.64 (m, 7H), 3.64 – 3.59 (m, 2H), 3.55 (t, *J* = 4.4 Hz, 2H), 3.50 – 3.44 (m, 2H), 2.58 – 2.40 (m, 3H), 2.30 (d, *J* = 13.0 Hz, 1H), 2.10 – 2.02 (m, 3H), 1.95 – 1.89 (m, 1H), 1.60 – 1.38 (m, 4H), 0.90 (d, *J* = 7.3 Hz, 3H). LCMS RT 4.53 min, ES(+) *m/z* 1008.5 $(M+H)^+$. HRMS (ES+) *m/z* calc. for $C_{53}H_{66}N_7O_{13}$ $(M+H)^+$: 1008.4719, found: 1008.4763.

**(*R*)-1-(3-((1-(4-(1,2,4,5-tetrazin-3-yl)phenyl)-3,19-dioxo-6,9,12,15-tetraoxa-2,18-diazaicosan-20-yl)oxy)phenyl)-3-(3,4-dimethoxyphenyl)propyl (*S*)-1-((*S*)-2-(3,4,5-trimethoxyphenyl)butanoyl)piperidine-2-carboxylate (FT2)**

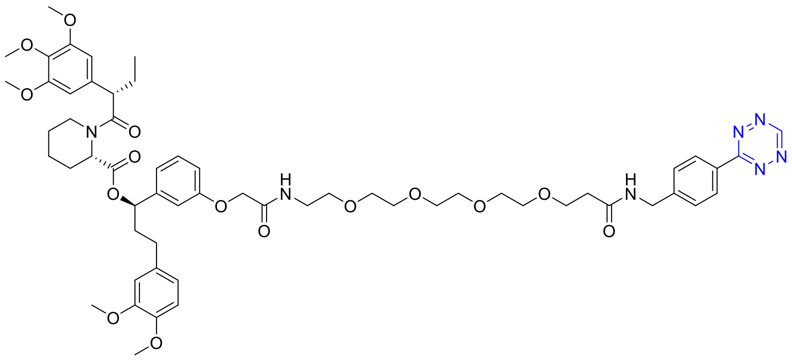

Compound **FT2** was synthesized according to the procedure reported for compound **FT1** using **2b** as the starting material. **FT2** was obtained as a pink solid, yield 38%. $R_f$ 0.3 (5% methanol in dichloromethane). $^1$H NMR (400 MHz, $CDCl_3$) δ 10.19 (s, 1H), 8.56 (d, *J* = 8.3 Hz, 2H), 7.52 (d, *J* = 8.4 Hz, 2H), 7.17 (t, *J* = 8.3 Hz, 1H), 7.09 (s, 1H), 6.76 (d, *J* = 6.7 Hz, 3H), 6.70 – 6.61

(m, 3H), 6.41 (d, *J* = 3.8 Hz, 2H), 5.66 – 5.57 (m, 1H), 5.44 (d, *J* = 6.0 Hz, 1H), 4.56 (d, *J* = 6.0 Hz, 2H), 4.46 (s, 2H), 3.86 – 3.82 (m, 10H), 3.80 – 3.66 (m, 18H), 3.64 – 3.59 (m, 4H), 3.51 (d, *J* = 5.6 Hz, 2H), 2.59 – 2.41 (m, 5H), 2.30 (d, *J* = 12.5 Hz, 1H), 2.06 (dq, *J* = 14.2, 7.8 Hz, 3H), 1.94 (dd, *J* = 15.0, 6.3 Hz, 1H), 1.70 – 1.58 (m, 4H), 0.89 (t, *J* = 7.3 Hz, 3H). LCMS RT 4.69 min, ES(+) *m/z* 1132.5 $(M+Na)^+$. HRMS (ES+) *m/z* calc. for $C_{58}H_{76}N_7O_{15}$ $(M+H)^+$: 1110.5399, found: 1110.5438.

**(*R*)-1-(3-((1-(4-(1,2,4,5-tetrazin-3-yl)phenyl)-3,25-dioxo-6,9,12,15,18,21-hexaoxa-2,24-diazahexacosan-26-yl)oxy)phenyl)-3-(3,4-dimethoxyphenyl)propyl (*S*)-1-((*S*)-2-(3,4,5-trimethoxyphenyl)butanoyl)piperidine-2-carboxylate (FT3)**

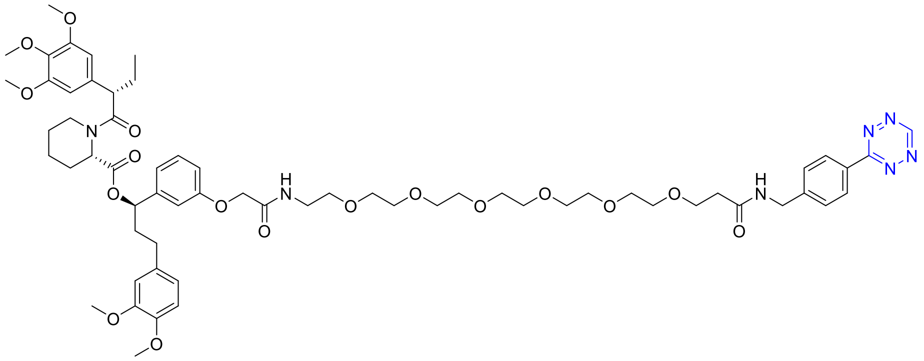

Compound **FT3** was synthesized according to the procedure reported for compound **FT1** using **2c** as the starting material. **FT3** was obtained as a pink solid, yield 88%. $R_f$ 0.4 (5% methanol in dichloromethane). $^1H$ NMR (400 MHz, $CDCl_3$) δ 10.19 (s, 1H), 8.56 (d, *J* = 8.4 Hz, 2H), 7.53 (d, *J* = 8.4 Hz, 2H), 7.32 – 7.27 (m, 1H), 7.17 (t, *J* = 8.3 Hz, 1H), 7.09 (s, 1H), 6.81 – 6.74 (m, 3H), 6.64 (d, *J* = 9.4 Hz, 3H), 6.40 (s, 2H), 5.61 (dd, *J* = 8.1, 5.4 Hz, 1H), 5.45 (d, *J* = 4.3 Hz, 1H), 4.57 (d, *J* = 6.1 Hz, 2H), 4.47 (d, *J* = 4.0 Hz, 2H), 3.87 – 3.81 (m, 10H), 3.79 – 3.74 (m, 12H), 3.67 – 3.62 (m, 10H), 3.56 – 3.52 (m, 10H), 2.62 – 2.40 (m, 5H), 2.30 (d, *J* = 13.0 Hz, 1H), 2.09 (d, *J* = 6.1 Hz, 3H), 1.96 – 1.87 (m, 1H), 1.68 – 1.34 (m, 4H), 0.93 – 0.85 (m, 3H). LCMS RT 4.72 min, ES(+) *m/z* 1198.6 $(M+H)^+$. HRMS (ES+) *m/z* calc. for $C_{62}H_{84}N_7O_{17}$ $(M+H)^+$: 1198.6924, found: 1198.5990.

The FKBP ligand-tetrazine probes were obtained through deprotection of compound **2a** and subsequent amide coupling with 6-chlorohexanoic acid **(Scheme 2)**.

**Scheme 2.** Synthesis of FKBP ligand-tetrazine probes[a]

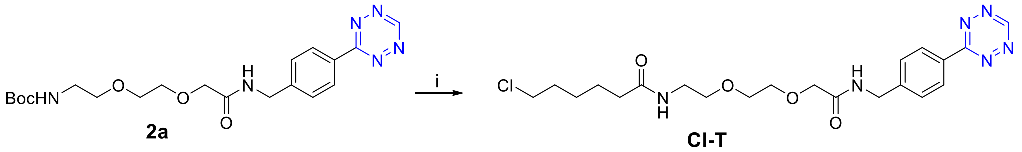

**2a** **Cl-T**

[a]Reagents and conditions. i) a. TFA, DCM, reflux, 1 h; b. 6-chlorohexanoic acid, EDCI, HOBt, DCM, rt, yield 54%.

##### *N*-(2-(2-(2-((4-(1,2,4,5-tetrazin-3-yl)benzyl)amino)-2-oxoethoxy)ethoxy)ethyl)-6-chlorohexanamide (Cl-T)

Compound **2a** (16 mg, 0.041 mmol) was added to a mixture of trifluoroacetic acid (2 mL) and dichloromethane (2 mL) and the reaction was refluxed for 1 h. The mixture was concentrated under reduced pressure and the crude was used in the next step without further purification. The deprotected product was added to a mixture in dichloromethane (5 mL) containing 6-chlorohexanoic acid (23 mg, 0.15 mmol), which was pre-activated by HATU (60 mg, 0.16 mmol) and triethylamine (50 uL, 0.35 mmol). The resulting mixture was stirred at room temperature for 18 h, then concentrated under reduced pressure. Purification by flash column chromatography (0-5% methanol in dichloromethane) afforded the title compound (11 mg, yield 58%) as a pink oil. $R_f$ 0.3 (5% methanol in dichloromethane). $^1$H NMR (400 MHz, $CDCl_3$) δ 10.22 (s, 1H), 8.60 (d, *J* = 8.3 Hz, 2H), 7.55 (d, *J* = 8.2 Hz, 2H), 6.11 (s, 1H), 5.98 (s, 1H), 3.69 (s, 2H), 3.57 – 3.51 (m, 8H), 3.48 – 3.39 (m, 4H), 2.23 – 2.17 (m, 2H), 1.86 – 1.64 (m, 6H). LCMS RT 3.53 min, ES(+) *m/z* 465.3 $(M+H)^+$. HRMS (ES+) *m/z* calc. for $C_{21}H_{30}ClN_6O_4$ $(M+H)^+$: 465.2017, found: 465.1867.

For the synthesis of Aurora targeting compounds, alisertib **3**[15] used as the Aurora ligand. Amide coupling with Boc-protected PEG-4 and PEG-5 based linkers and subsequent Boc-deprotection afforded amine intermediates **5a** and **5b**. A final amide coupling step using 2-(4-(1,2,4,5-tetrazin-3-yl)phenyl)acetic acid provided final compounds **6a** and **6b (Scheme 2).**

**Scheme 2.** Synthesis of Aurora targeting compounds[a]

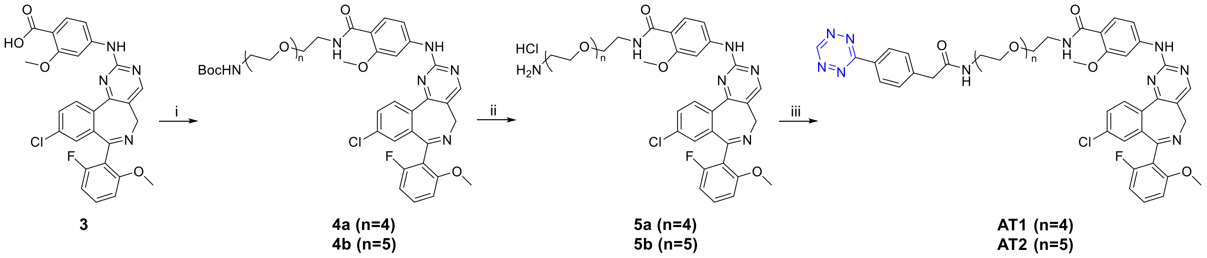

[a]Reagents and conditions. i. tert-butyl (14-amino-3,6,9,12-tetraoxatetradecyl)carbamate or tert-butyl (17-amino-3,6,9,12,15-pentaoxaheptadecyl)carbamate, T3P, DIPEA, DMF, rt, 18 h, yield 24-54%; ii. HCl in dioxane 4M, dioxane, rt, 2 h, yield 90%-quant.; iii. 2-(4-(1,2,4,5-tetrazin-3-yl)phenyl)acetic acid, T3P, DIPEA, DMF, rt, 18 h, yield 22-58%.

##### tert-butyl (1-(4-((9-chloro-7-(2-fluoro-6-methoxyphenyl)-5*H*-benzo[*c*]pyrimido[4,5-*e*]azepin-2-yl)amino)-2-methoxyphenyl)-1-oxo-5,8,11,14-tetraoxa-2-azahexadecan-16-yl)carbamate (4a)

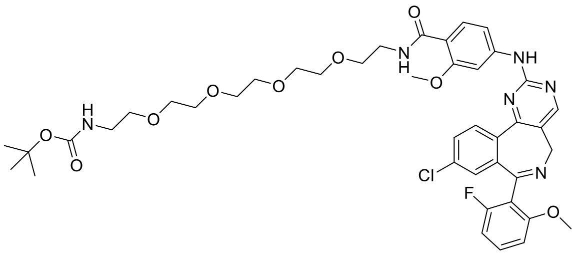

To a solution of 4-((9-chloro-7-(2-fluoro-6-methoxyphenyl)-5*H*-benzo[*c*]pyrimido[4,5-*e*]azepin-2-yl)amino)-2-methoxybenzoic acid **(3)** (34 mg, 0.066 mmol) in anhydrous DMF (2 mL) under nitrogen, was added tert-butyl (14-amino-3,6,9,12-tetraoxatetradecyl)carbamate (22 mg, 0.066 mmol), 1-propanephosphonic anhydride 50% in DMF (46 uL, 0.079 mmol) and *N,N*-diisopropylethylamine (46 uL, 0.26 mmol), and the reaction was stirred at room temperature for 18 h. The mixture was diluted with ethyl acetate and was washed with LiCl 5% solution (x2) and brine. The organic layer was collected, dried over sodium sulfate, filtered and concentrated under reduced pressure. Purification by flash column chromatography (0-5% methanol in ethyl acetate) afforded the title compound (13 mg, yield 24%) as a colourless oil which slowly crystallised. $R_f$ 0.4 (5% methanol in dichloromethane). $^1$H NMR (400 MHz, $CDCl_3$) δ 8.52 (s, 1H), 8.25 – 8.15 (m, 3H), 7.93 (d, *J* = 2.0 Hz, 1H), 7.62 (s, 1H), 7.56 (dd, *J* = 8.5, 2.1 Hz, 1H), 7.35 – 7.27 (m, 2H), 7.10 (d, *J* = 8.5 Hz, 1H), 6.95 – 6.46 (m, 2H), 5.12 (s, 1H), 4.88 (s, 1H), 3.98 (s, 6H), 3.69 – 3.58 (m, 16H), 3.51 (t, *J* = 5.0 Hz, 2H), 3.37 – 3.23 (m, 3H), 1.43 (s, 9H). LCMS RT 4.58 min, ES(+) *m/z* 837.4 $(M+H)^+$.

**tert-butyl (1-(4-((9-chloro-7-(2-fluoro-6-methoxyphenyl)-5*H*-benzo[*c*]pyrimido[4,5-*e*]azepin-2-yl)amino)-2-methoxyphenyl)-1-oxo-5,8,11,14,17-pentaoxa-2-azanonadecan-19-yl)carbamate (4b)**

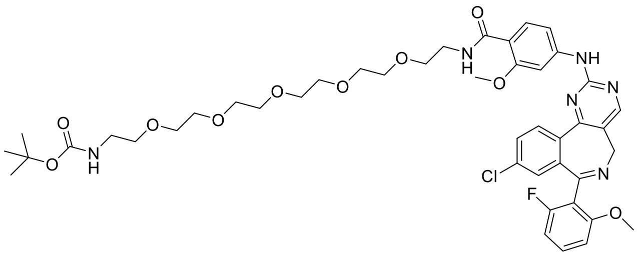

Compound **4b** was synthesized according to the procedure reported for compound **4a** using tert-butyl (17-amino-3,6,9,12,15-pentaoxaheptadecyl)carbamate. **4b** was obtained as a colourless oil which slowly crystallised, yield 54%. $R_f$ 0.5 (10% methanol in dichloromethane). $^1$H NMR (400 MHz, $CDCl_3$) δ 8.51 (s, 1H), 8.25 – 8.11 (m, 3H), 7.91 (d, *J* = 2.1 Hz, 1H), 7.79 – 7.66 (m, 1H), 7.55 (m, 1H), 7.33 – 7.27 (m, 2H), 7.18 – 7.09 (m, 1H), 6.95 – 6.39 (m, 2H), 5.18 (s, 1H), 4.87 (s, 1H), 3.98 (s, 6H), 3.69 – 3.57 (m, 20H), 3.51 (t, *J* = 5.2 Hz, 2H), 3.36 – 3.20 (m, 3H), 1.42 (s, 9H). LCMS RT 5.30 min, ES(+) *m/z* 881.4 $(M+H)^+$.

***N*-(14-amino-3,6,9,12-tetraoxatetradecyl)-4-((9-chloro-7-(2-fluoro-6-methoxyphenyl)-5*H*-benzo[*c*]pyrimido[4,5-e]azepin-2-yl)amino)-2-methoxybenzamide hydrochloride (5a)**

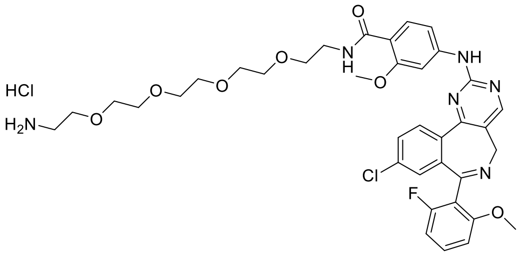

To a solution of compound **4a** (13 mg, 0.016 mmol) in dioxane (1 mL), was added HCl in dioxane 4 M (1 mL) dropwise at 0 °C and the reaction was stirred at room temperature for 2 h. The mixture was concentrated under reduced pressure and the residue was triturated with diethyl ether to afford the title compound (12 mg, yield quant.) as a light yellow solid. $^1$H NMR

(400 MHz, DMSO-$d_6$) δ 10.21 (s, 1H), 8.72 (s, 1H), 8.30 (d, *J* = 8.5 Hz, 1H), 8.19 (t, *J* = 5.5 Hz, 1H), 7.97 (s, 1H), 7.94 – 7.66 (m, 5H), 7.47 – 7.38 (m, 2H), 7.23 (s, 1H), 7.10 – 6.74 (m, 2H), 4.87 (s, 1H), 3.93 (s, 6H), 3.63 – 3.53 (m, 19H), 3.00 – 2.90 (m, 2H). LCMS RT 3.71 min, ES(+) *m/z* 737.4 $(M+H)^+$.

***N*-(17-amino-3,6,9,12,15-pentaoxaheptadecyl)-4-((9-chloro-7-(2-fluoro-6-methoxyphenyl)-5*H*-benzo[*c*]pyrimido[4,5-*e*]azepin-2-yl)amino)-2-methoxybenzamide hydrochloride (5b)**

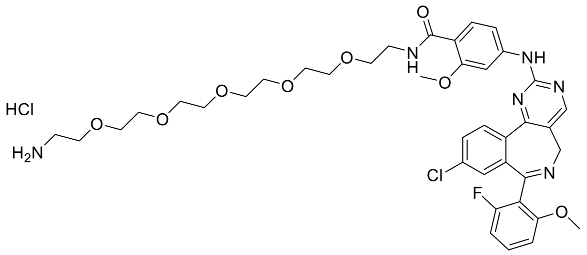

Compound **5b** was synthesized according to the procedure reported for compound **5a** using **4b** as the starting material. **5b** was obtained as a light yellow solid, yield 90%. $^1$H NMR (400 MHz, DMSO-$d_6$) δ 10.23 (s, 1H), 8.73 (s, 1H), 8.32 (d, *J* = 8.5 Hz, 1H), 8.18 (t, *J* = 5.4 Hz, 1H), 7.98 (s, 1H), 7.95 – 7.74 (m, 5H), 7.48 – 7.38 (m, 2H), 7.25 (s, 1H), 7.00 – 6.77 (m, 2H), 4.88 (s, 1H), 3.93 (s, 4H), 3.60 – 3.44 (m, 25H), 2.99 – 2.90 (m, 2H). LCMS RT 4.53 min, ES(+) *m/z* 781.4 $(M+H)^+$.

***N*-(1-(4-(1,2,4,5-tetrazin-3-yl)phenyl)-2-oxo-6,9,12,15-tetraoxa-3-azaheptadecan-17-yl)-4-((9-chloro-7-(2-fluoro-6-methoxyphenyl)-5*H*-benzo[*c*]pyrimido[4,5-*e*]azepin-2-yl)amino)-2-methoxybenzamide (AT1)**

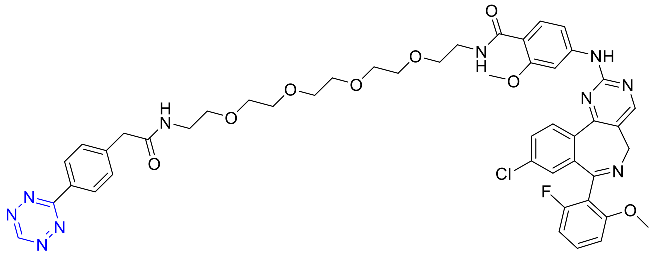

To a solution of compound **5a** (10 mg, 0.013 mmol) in anhydrous DMF (2 mL) under nitrogen, was added 2-(4-(1,2,4,5-tetrazin-3-yl)phenyl)acetic acid (2.8 mg, 0.013 mmol), 1-propanephosphonic anhydride 50% in DMF (9 uL, 0.016 mmol) and *N,N*-diisopropylethylamine (9 uL, 0.052 mmol), and the reaction was stirred at room temperature for 18 h. The mixture was diluted with ethyl acetate and was washed with LiCl 5% solution (x2) and brine. The organic layer was collected, dried over sodium sulfate, filtered and concentrated under reduced pressure. Purification by flash column chromatography (0-5% methanol in dichloromethane) afforded the title compound (7.0 mg, yield 58%) as a pink solid. $R_f$ 0.4 (5% methanol in dichloromethane). $^1$H NMR (400 MHz, DMSO-$d_6$) δ 10.56 (s, 1H), 10.20 (s, 1H), 8.72 (s, 1H), 8.42 (d, *J* = 8.0 Hz, 2H), 8.30 (d, *J* = 8.5 Hz, 1H), 8.25 (d, *J* = 5.5 Hz, 1H), 8.17 (t, *J* = 5.4 Hz, 1H), 7.97 (s, 1H), 7.85 (d, *J* = 8.6 Hz, 1H), 7.81 (dd, *J* = 8.5, 2.0 Hz, 1H), 7.54 (d, *J* = 8.1 Hz, 2H), 7.46 – 7.37 (m, 2H), 7.21 (s, 1H), 7.02 – 6.71 (br, 2H), 4.99-4.75 (br, 1H), 3.92 (s, 5H), 3.64 – 3.37 (m, 22H), 3.22 (q, *J* = 5.7 Hz, 2H). LCMS RT 4.80 min, ES(+)

$m/z$ 935.4 $(M+H)^+$. HRMS (ES+) $m/z$ calc. for $C_{47}H_{49}ClFN_{10}O_8$ $(M+H)^+$: 935.3407, found: 935.3439.

**N-(1-(4-(1,2,4,5-tetrazin-3-yl)phenyl)-2-oxo-6,9,12,15,18-pentaoxa-3-azaicosan-20-yl)-4-((9-chloro-7-(2-fluoro-6-methoxyphenyl)-5*H*-benzo[*c*]pyrimido[4,5-*e*]azepin-2-yl)amino)-2-methoxybenzamide (AT2)**

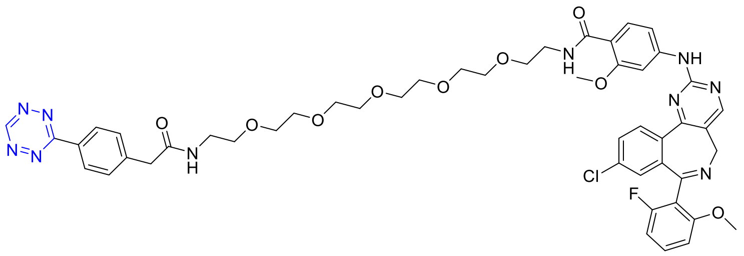

Compound **AT2** was synthesized according to the procedure reported for compound **AT2** using **5b** as the starting material. **AT2** was obtained as a pink solid, yield 22%. $R_f$ 0.4 (5% methanol in dichloromethane). $^1$H NMR (400 MHz, DMSO-*d6*) δ 10.56 (s, 1H), 10.20 (s, 1H), 8.72 (s, 1H), 8.42 (d, *J* = 8.1 Hz, 2H), 8.30 (d, *J* = 8.5 Hz, 1H), 8.25 (t, *J* = 5.4 Hz, 1H), 8.17 (t, *J* = 5.4 Hz, 1H), 7.98 (s, 1H), 7.85 (d, *J* = 8.7 Hz, 1H), 7.81 (dd, *J* = 8.4, 2.2 Hz, 1H), 7.54 (d, *J* = 8.1 Hz, 2H), 7.47 - 7.36 (m, 2H), 7.22 (s, 1H), 7.07 - 6.69 (br, 2H), 5.01 - 4.73 (br, 1H), 3.92 (s, 5H), 3.61 - 3.38 (m, 26H), 3.22 (q, *J* = 5.7 Hz, 2H). LCMS RT 5.13 min, ES(+) $m/z$ 979.4 $(M+H)^+$. HRMS (ES+) $m/z$ calc. for $C_{49}H_{53}ClFN_{10}O_9$ $(M+H)^+$: 979.3670, found: 979.3696.

### Final compounds – Characterization data

#### Compound **FT1**

1. $^1$H-NMR

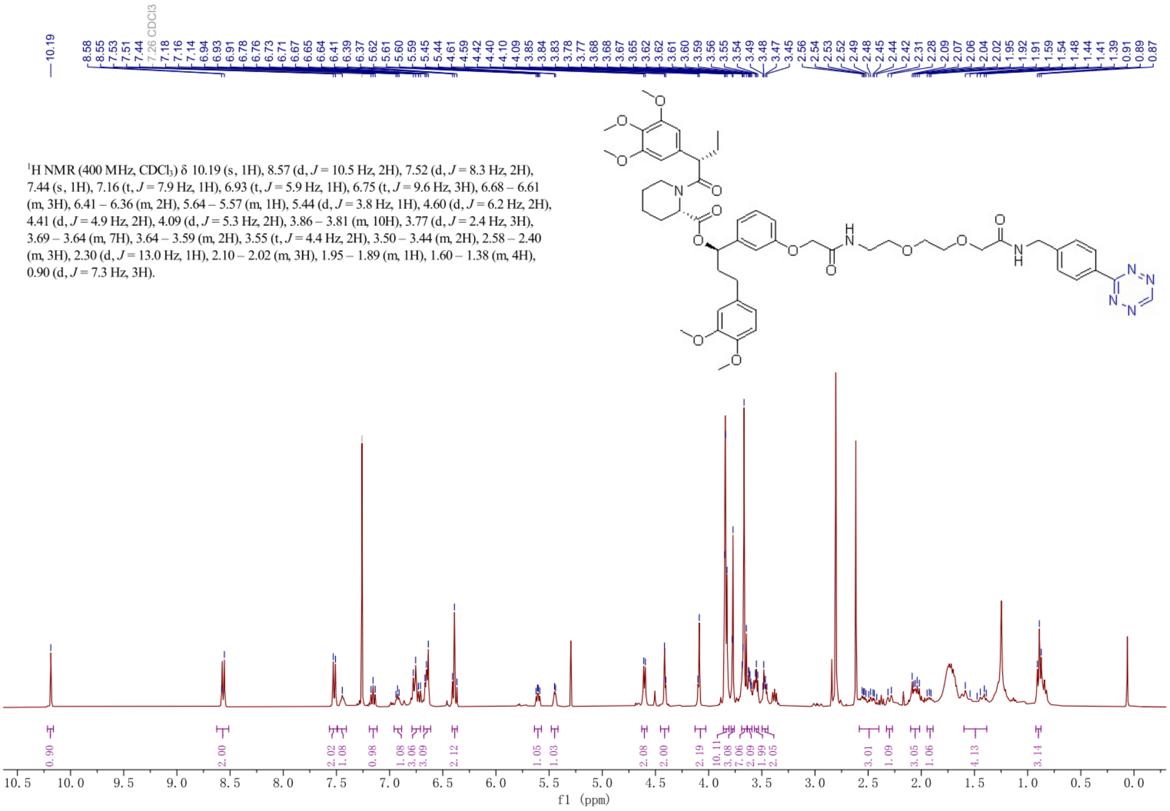

2. HRMS

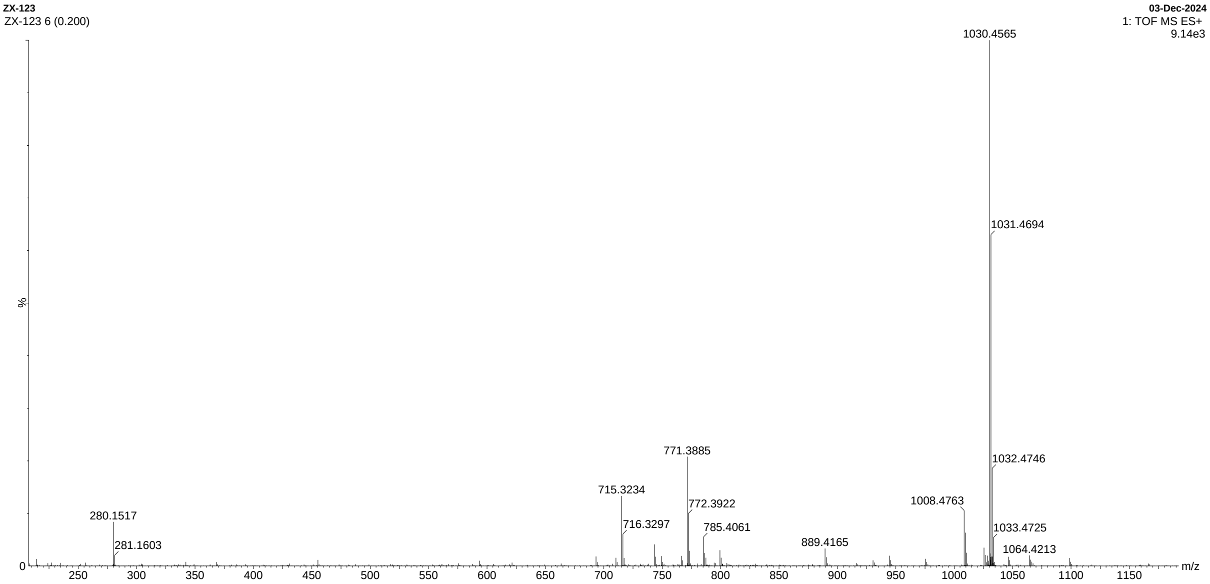

[M+H]+ m/z 1008.4763; [M+Na]+ m/z 1030.4565

Compound **FT2**

1. $^{1}$H-NMR

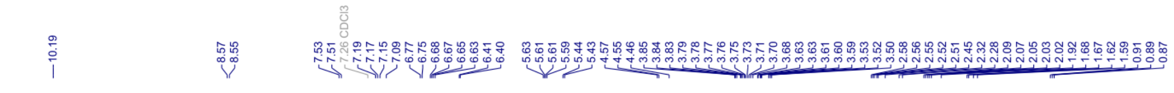

[1]H NMR (400 MHz, CDCl$_3$) δ 10.19 (s, 1H), 8.56 (d, *J* = 8.3 Hz, 2H), 7.52 (d, *J* = 8.4 Hz, 2H), 7.17 (t, *J* = 8.3 Hz, 1H), 7.09 (s, 1H), 6.76 (d, *J* = 6.7 Hz, 3H), 6.70 – 6.61 (m, 3H), 6.41 (d, *J* = 3.8 Hz, 2H), 5.66 – 5.57 (m, 1H), 5.44 (d, *J* = 6.0 Hz, 1H), 4.56 (d, *J* = 6.0 Hz, 2H), 4.46 (s, 2H), 3.86 – 3.82 (m, 10H), 3.80 – 3.66 (m, 18H), 3.64 – 3.59 (m, 4H), 3.51 (d, *J* = 5.6 Hz, 2H), 2.59 – 2.41 (m, 5H), 2.30 (d, *J* = 12.5 Hz, 1H), 2.06 (dq, *J* = 14.2, 7.8 Hz, 3H), 1.94 (dd, *J* = 15.0, 6.3 Hz, 1H), 1.70 – 1.58 (m, 4H), 0.89 (t, *J* = 7.3 Hz, 3H).

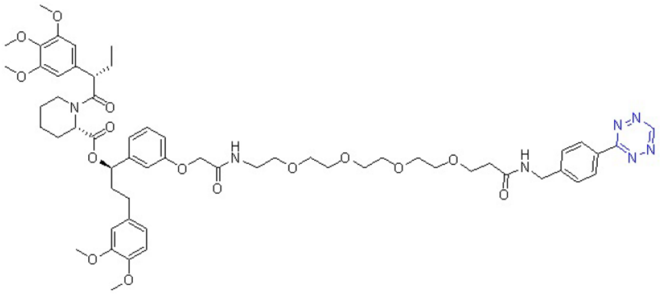

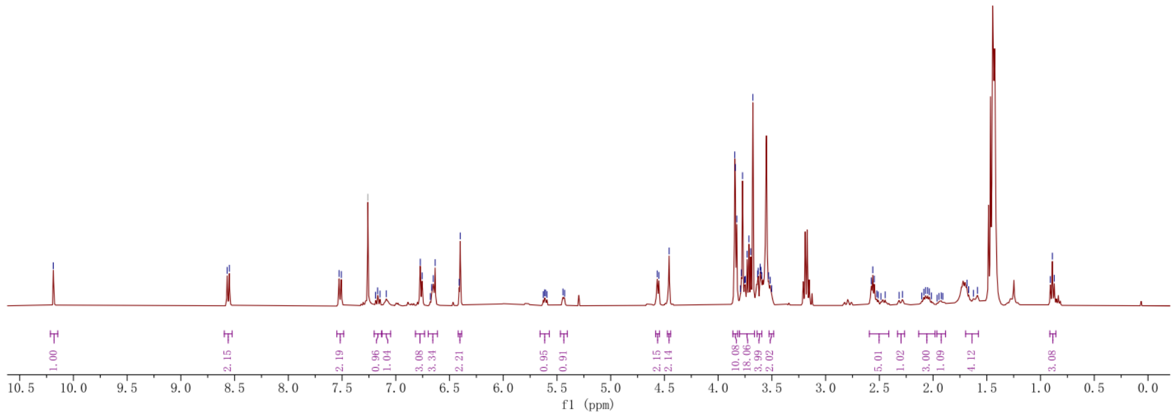

2. HRMS

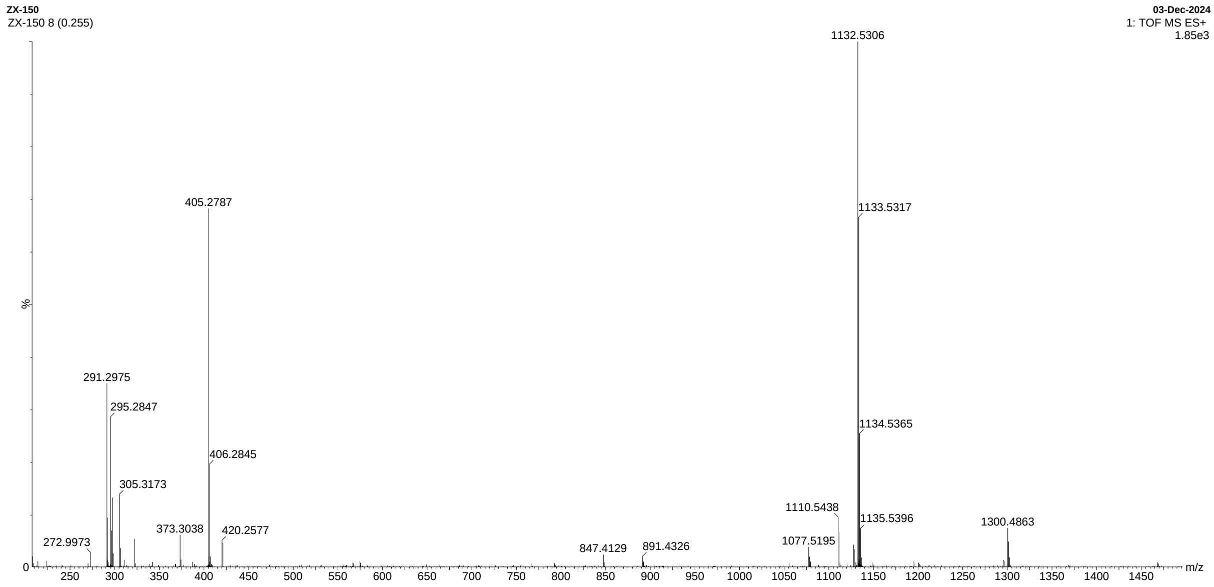

[M+H]$^{+}$ *m/z* 1110.5438; [M+Na]$^{+}$ *m/z* 1132.5306

Compound **FT3**

1. $^{1}$H-NMR

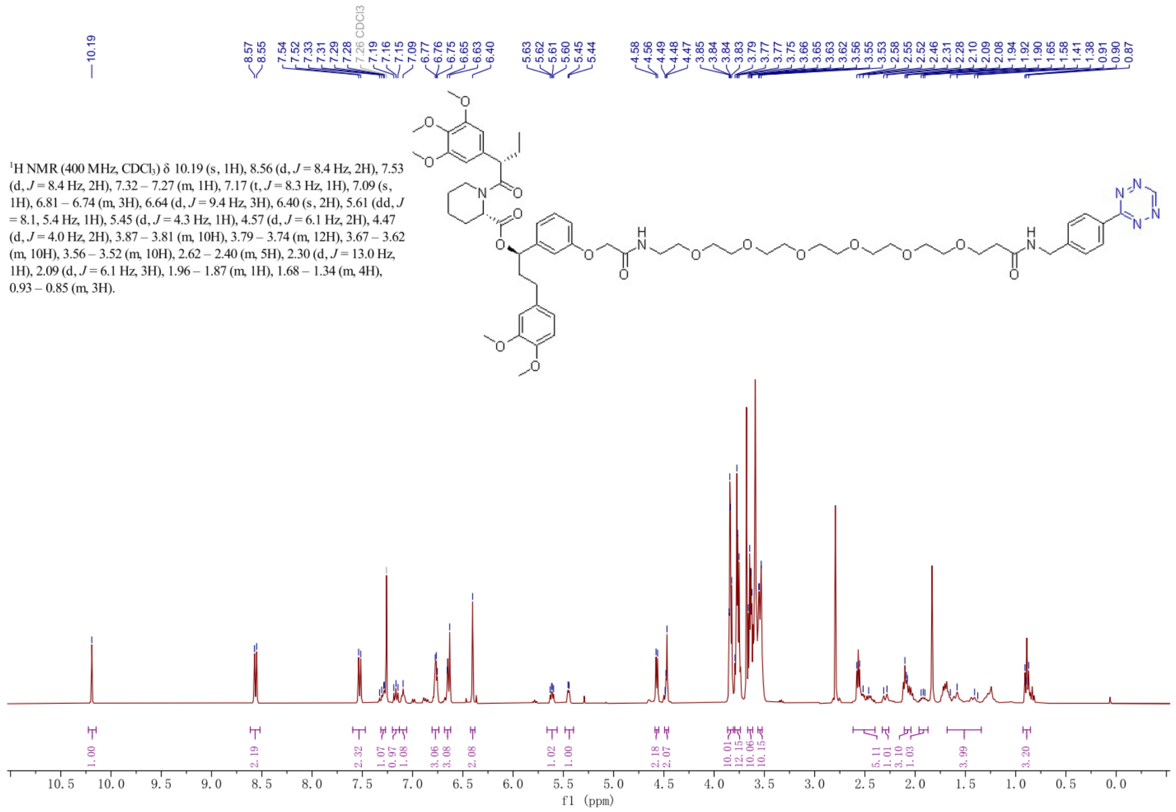

2. HRMS

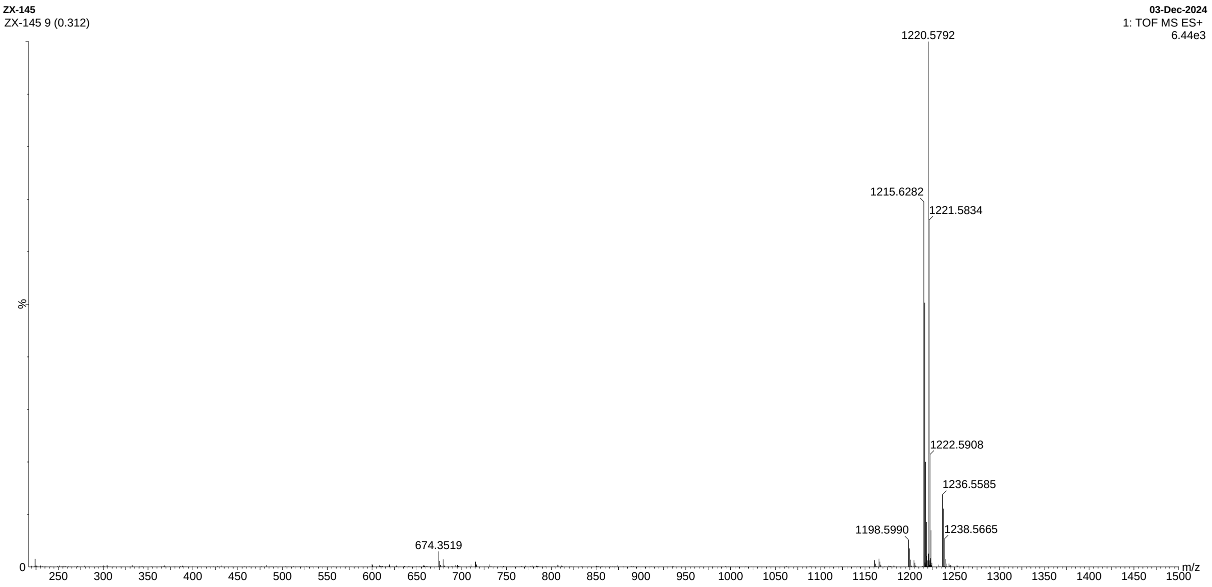

$[M+H]^+$ *m/z* 1198.5990; $[M+NH_4]^+$ *m/z* 1215.6282; $[M+Na]^+$ *m/z* 1220.5792

Compound **Cl-T**

1. $^1$H-NMR

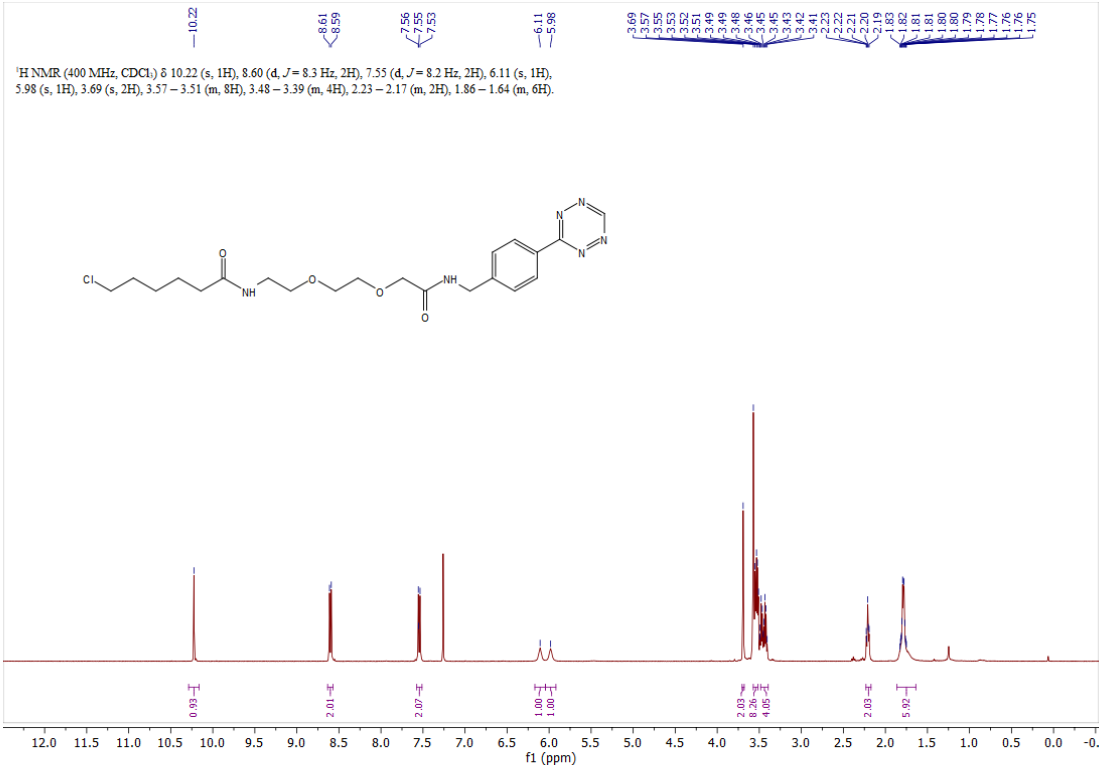

2. HRMS

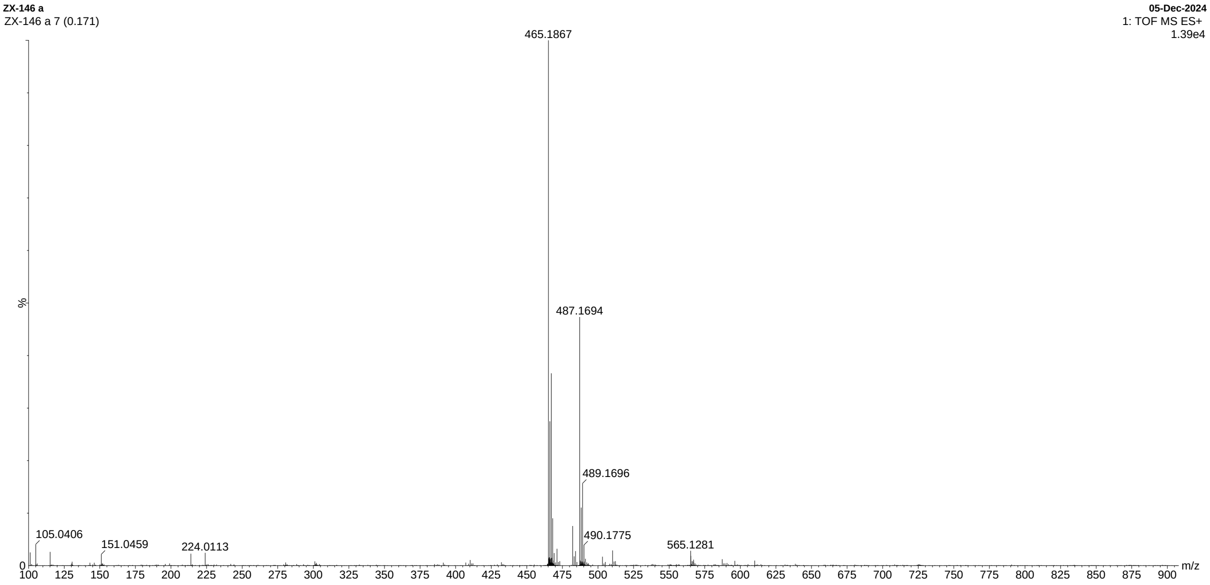

[M+H]$^+$ *m/z* 465.1867; [M+Na]$^+$ *m/z* 487.1694

Compound **AT1**

1. $^{1}$H-NMR

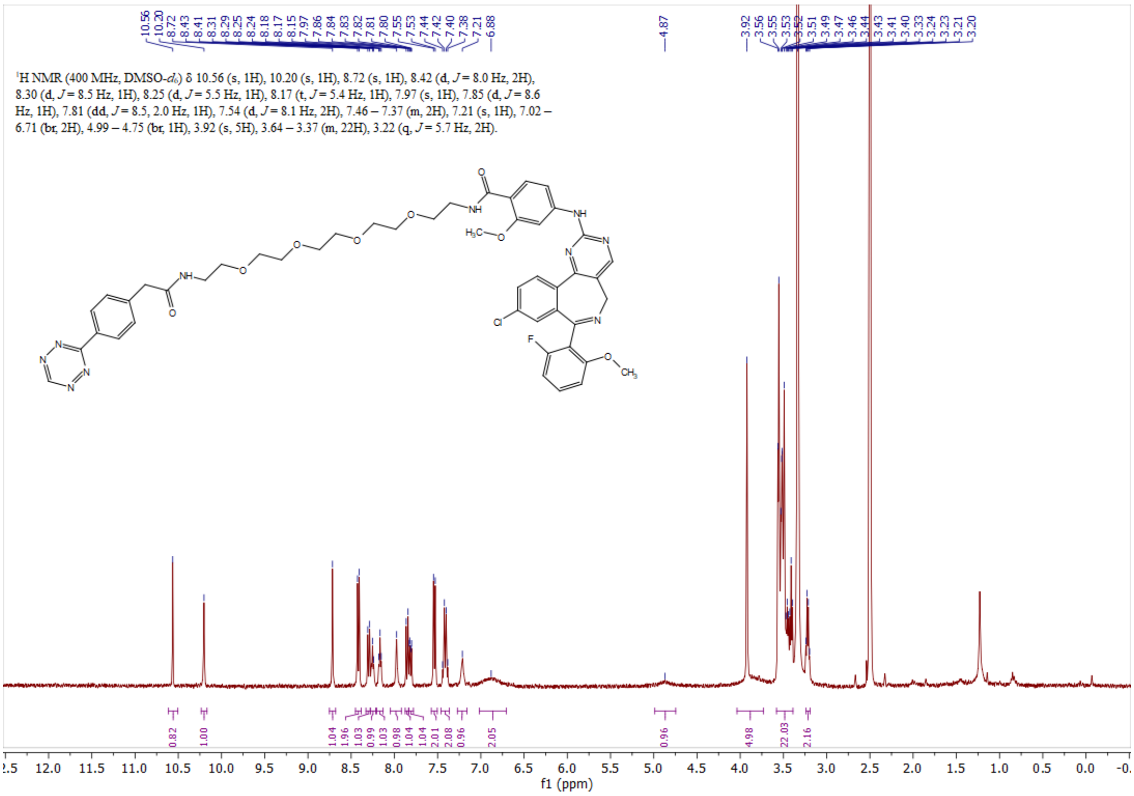

2. HRMS

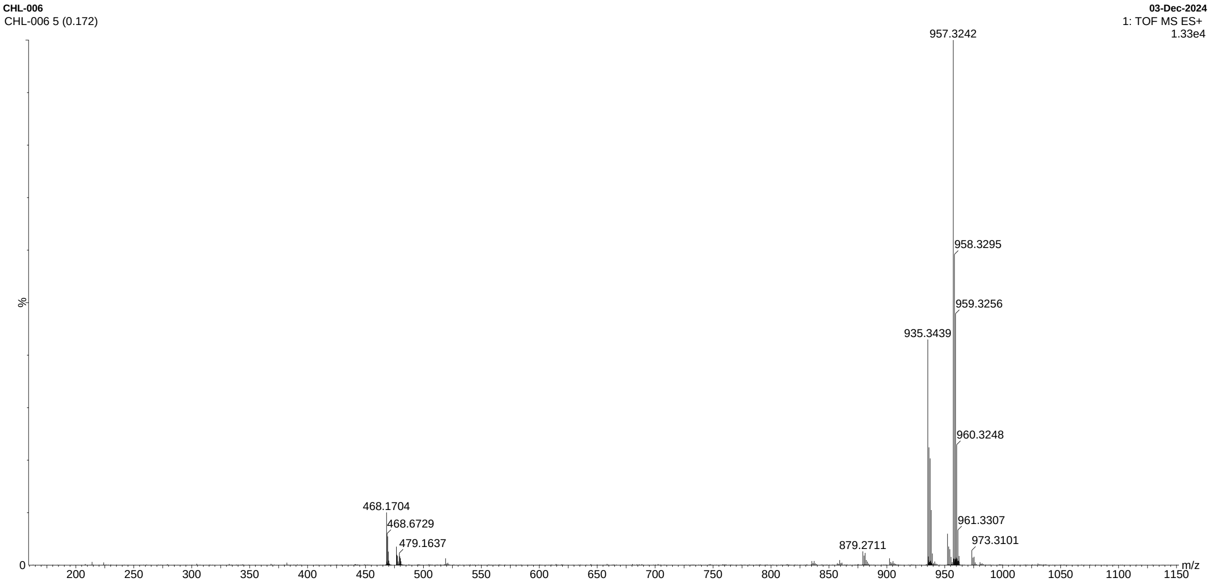

[M+H]$^{+}$ *m/z* 935.3439; [M+Na]$^{+}$ *m/z* 957.3242

Compound **AT2**

1. [1]H-NMR

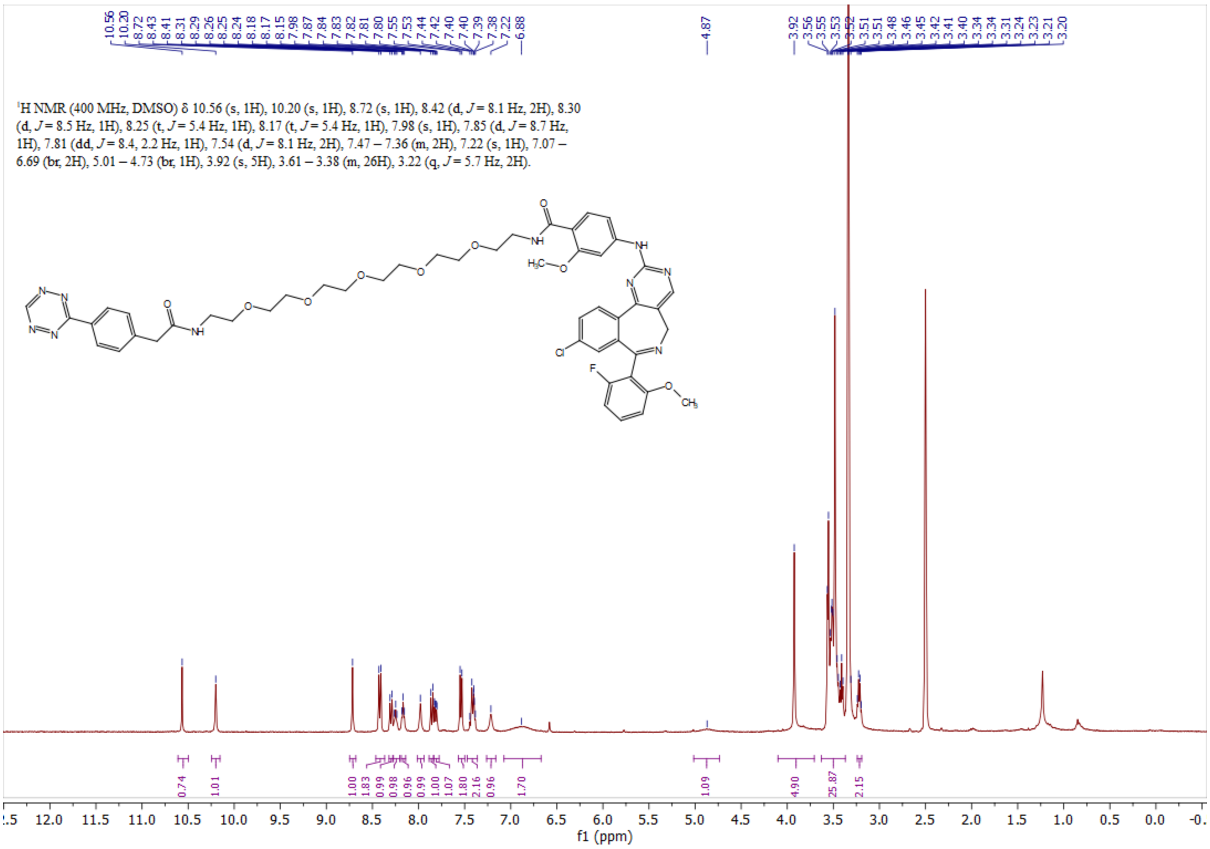

2. HRMS

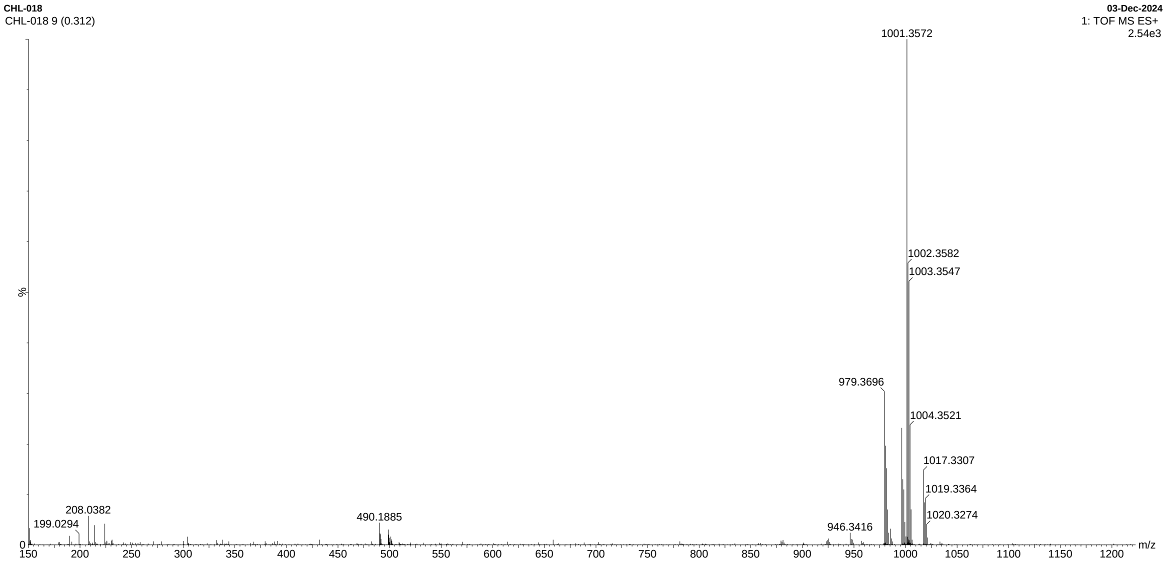

$[M+H]^+$ *m/z* 979.3696; $[M+Na]^+$ *m/z* 1001.3572

#### Reference

(1) Nabet, B.; Ferguson, F. M.; Seong, B. K. A.; Kuljanin, M.; Leggett, A. L.; Mohardt, M. L.; Robichaud, A.; Conway, A. S.; Buckley, D. L.; Mancias, J. D.; et al. Rapid and Direct Control of Target Protein Levels with VHL-Recruiting DTAG Molecules. *Nat. Commun.* **2020**, *11* (1).

(2) Nabet, B.; Roberts, J. M.; Buckley, D. L.; Paulk, J.; Dastjerdi, S.; Yang, A.; Leggett, A. L.; Erb, M. A.; Lawlor, M. A.; Souza, A.; et al. The DTAG System for Immediate and Target-Specific Protein Degradation. *Nat. Chem. Biol.* **2018**, *14* (5), 431–441.

(3) Liu, Y.; Yang, J.; Wang, T.; Luo, M.; Chen, Y.; Chen, C.; Ronai, Z.; Zhou, Y.; Ruppin, E.; Han, L. Expanding PROTACtable Genome Universe of E3 Ligases. *Nat. Commun.* **2023**, *14* (1).

(4) Boatner, L. M.; Palafox, M. F.; Schweppe, D. K.; Backus, K. M. CysDB: A Human Cysteine Database Based on Experimental Quantitative Chemoproteomics. *Cell Chem. Biol.* **2023**, *30* (6), 683-698.e3.

(5) Kuljanin, M.; Mitchell, D. C.; Schweppe, D. K.; Gikandi, A. S.; Nusinow, D. P.; Bulloch, N. J.; Vinogradova, E. V.; Wilson, D. L.; Kool, E. T.; Mancias, J. D.; et al. Reimagining High-Throughput Profiling of Reactive Cysteines for Cell-Based Screening of Large Electrophile Libraries. *Nat. Biotechnol.* **2021**, *39* (5), 630–641.

(6) Takahashi, M.; Chong, H. B.; Zhang, S.; Yang, T. Y.; Lazarov, M. J.; Harry, S.; Maynard, M.; Hilbert, B.; White, R. D.; Murrey, H. E.; et al. DrugMap: A Quantitative Pan-Cancer Analysis of Cysteine Ligandability. *Cell* **2024**, *187* (10), 2536-2556.e30.

(7) Ignatov, M.; Jindal, A.; Kotelnikov, S.; Beglov, D.; Posternak, G.; Tang, X.; Maisonneuve, P.; Poda, G.; Batey, R. A.; Sicheri, F.; et al. High Accuracy Prediction of PROTAC Complex Structures. *J. Am. Chem. Soc.* **2023**, *145*, 7123–7135.

(8) Sergei Kotelnikov, Andrey Alekseenko, Cong Liu, Mikhail Ignatov, Dzmitry Padhorny, Emiliano Brini, Mark Lukin, Evangelos Coutsias, K. A. D. & D. K. Sampling and Refinement Protocols for Template-Based Macrocycle Docking: 2018 D3R Grand Challenge 4. *J. Comput. Aided. Mol. Des.* **2020**, *34*, 179–189.

(9) Alekseenko, A.; Kotelnikov, S.; Ignatov, M.; Egbert, M.; Kholodov, Y.; Vajda, S.; Kozakov, D. ClusPro LigTBM: Automated Template-Based Small Molecule Docking. *J. Mol. Biol.* **2020**, *432*, 3404–3410.

(10) Kotelnikov, S.; Ashizawa, R.; Popov, K. I.; Khan, O.; Ignatov, M.; Li, S. X.; Hassan, M.; Coutsias, E. A.; Poda, G.; Padhorny, D.; et al. Accurate Ligand–Protein Docking in CASP15 Using the ClusPro LigTBM Server. *Proteins Struct. Funct. Bioinforma.* **2023**, *91* (12), 1822–1828.

(11) Kozakov, D.; Brenke, R.; Comeau, S. R.; Vajda, S. PIPER: An FFT-Based Protein Docking Program with Pairwise Potentials. *Proteins* **2006**, *65*, 392–406.

(12) Riniker, S.; Landrum, G. A. Better Informed Distance Geometry: Using What We Know To Improve Conformation Generation. *J. Chem. Inf. Model.* **2015**, *55*, 2562–2574.

(13) Kozakov, D.; Hall, D. R.; Xia, B.; Porter, K. A.; Padhorny, D.; Yueh, C.; Beglov, D.; Vajda, S. The ClusPro Web Server for Protein – Protein Docking. *Nat. Protoc.* **2017**, *12* (2), 255–278.

(14) Case, D. A.; Cheatham, T. E.; Darden, T.; Gohlke, H.; Luo, R.; Merz, K. M.; Onufriev, A.; Simmerling, C.; R, B. W. and; Woods, J. The Amber Biomolecular Simulation Programs. *J. Comput. Chem.* **2005**, *26*, 1668–1688.

(15) Sells, T. B.; Chau, R.; Ecsedy, J. A.; Gershman, R. E.; Hoar, K.; Huck, J.; Janowick, D. A.; Kadambi, V. J.; Leroy, P. J.; Stirling, M.; et al. MLN8054 and Alisertib (MLN8237): Discovery of Selective Oral Aurora A Inhibitors. *ACS Med. Chem. Lett.* **2015**, *6* (6), 630–634.
